# Cholinergic signaling differentially regulates a vocal motor pallial nucleus to stabilize adult birdsong

**DOI:** 10.1101/2024.09.03.610982

**Authors:** Ning Xu, Yutao Zhang, Yalun Sun, Xueqing Song, Yangyang Cao, Xinqi Yang, Yuejun Zhou, Wei Meng

## Abstract

Cholinergic modulation plays an important role in motor skill performance, including vocal control. In songbirds, song motor nucleus RA simultaneously receives inputs from song nuclei HVC and LMAN, and then its projection neurons (RAPNs) generate song motor control output. Using electrophysiological and pharmacological methods, we found that cholinergic signaling can enhance song stability by reducing HVC-RAPN excitatory synaptic transmission in adult male zebra finches, mediated by mAChRs. Although nAChRs are not effective overall, cholinergic signaling can also decrease LMAN-RAPN excitatory synaptic transmission induced by electrical stimulation via nAChRs, suggesting the potential role of cholinergic regulation in song behavior through LMAN-RA pathway. On the contrary, in adult female zebra finches, only LMAN-RAPN synaptic transmission was reduced by cholinergic signaling via mAChRs. The role of differential cholinergic regulation of song premotor circuits in songbirds’ singing provides insights into the neural control mechanism of motor skill performance.

## INTRODUCTION

The long-term maintenance of learned motor skills, as well as their short-term plasticity and acute modulation, play a fundamental role throughout an individual’s life, such as writing, cycling, playing the piano, dancing, and speaking^1^. Learned motor skills are typically acquired gradually through multiple learning sessions until performance reaches stability^2^, within which cholinergic signaling assumes a pivotal role^3, 4^. However, the process and mechanism of cholinergic regulation on the long-term maintenance of learned motor skills and their short-term plasticity and acute modulation remain unclear.

Similar to human speech, songbird singing is a rare vocal learning behavior in the animal kingdom, and it is also a complex learned motor skill^5^. Juvenile songbirds optimize the matching degree between their own singing and the instructional song template through repetitive practice and self-correction relying on auditory feedback, and their songs progressively stabilize during the course of attaining adulthood^6^. Two well-defined neural pathways in songbird brain, namely vocal motor pathway (VMP) responsible for vocal production and anterior forebrain pathway (AFP) responsible for juvenile song learning and adult song plasticity, coordinately control singing^7, 8^. VMP consists of the song premotor nucleus HVC (proper name) and the song motor pallial nucleus RA (robust nucleus of the arcopallium), and brainstem motor nuclei; AFP is a cortex-basal ganglia-thalamus circuit in which the cortex nucleus, the lateral magnocellular nucleus of the anterior nidopallium (LMAN), efferently projects to RA of VMP, and the basal ganglia receives afferent projection from HVC of VMP^9^.

As an analogous structure of human laryngeal motor cortex, RA serves as the convergence nucleus within song premotor circuits, and concurrently receives glutamatergic projections from its upstream premotor nucleus HVC and LMAN of AFP (refer to Figure 1A) ^10, 11^. Meanwhile, RA receives cholinergic projections from the ventral pallidum (VP) in basal forebrain^12, 13^. During the critical period of song learning, an elevation of acetylcholine (ACh) concentration within RA was observed in male zebra finches^14^. Although the expression of both muscarinic ACh receptors (mAChRs) and nicotinic ACh receptors (nAChRs) are lower inside the RA of male zebra finches compared to the surround, these two types of ACh receptors exist in RA at every developmental stage^15, 16^. The combined infusion of mAChR and nAChR antagonists into RA leads to abnormal song development during the critical period^17^. RA consists of two types of neurons, including RA interneurons and RA projection neurons (RAPNs) that are uniquely associated with the acoustic properties of song subsyllables and input the information to brainstem motor nuclei^18, 19, 20, 21^. Our previous work showed that cholinergic signaling primarily modulates RAPNs’ electrophysiological activities in adult male zebra finches via mAChRs, but not nAChRs^22^. These studies indicate that RA is a crucial target of cholinergic modulation. Nonetheless, the essential mechanism by which cholinergic signaling governs RA to impact song production remains unknown. To clarify this issue, it is indispensable to understand the mechanism through which cholinergic signaling regulates the song premotor circuits with RA as the core.

**Figure 1.**
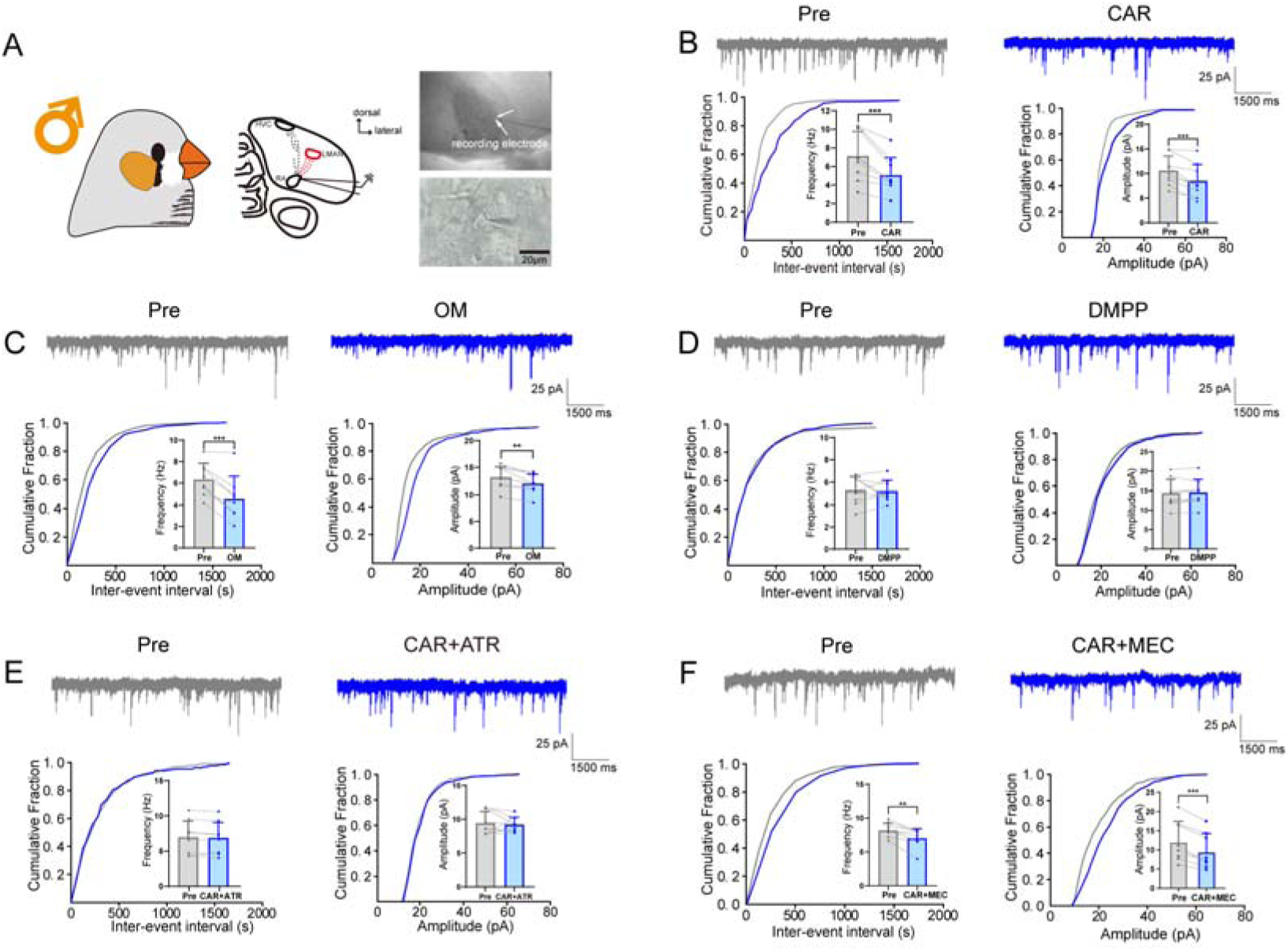
Cholinergic modulation of RAPNs’ mEPSCs in adult male zebra finches. (A) Experiment schematic. The pictures show in vitro whole-cell patch clamp recording of RAPNs’ mEPSCs in adult males. (B) Top, example traces of adult male RAPNs’ mEPSCs in Pre and CAR; Bottom, quantification of mEPSCs’ frequency (left) and amplitude (right) in Pre and CAR, n = 9. (C-F) Similar to (B). OM (C), n = 8; DMPP (D), n = 9; CAR+ATR (E), n = 9; CAR+MEC (F), n = 8. The variable n refers to the number of cells. ** *p* ≤ 0.01, *** *p* ≤ 0.001, paired *t*-test.

In this study, we initially employed in vitro patch-clamp whole-cell recording combined with pharmacological approaches to separately examine the roles of two cholinergic receptors, mAChRs and nAChRs, in cholinergic modulation of RAPNs’ excitatory synaptic afferents in adult male zebra finches. Given that the song premotor signals input to RA are respectively derived from HVC and LMAN, we conducted a further investigation into the influences of different cholinergic receptors on HVC-RAPN and LMAN-RAPN excitatory synaptic transmission. Subsequently, we used in vivo targeted pharmacological manipulation combined with song analysis to detect the effects of different cholinergic receptors within RA on birdsongs. Moreover, considering that zebra finches manifest typical sexual dimorphism, which only males sing and females never do^10^, we also explored whether the cholinergic modulation of the song premotor circuits are similar in adult male and female zebra finches. Overall, our findings delineate a sex difference mechanism by which cholinergic signaling differentially regulates song premotor circuits via mAChRs and nAChRs. This kind of differential regulation of song premotor circuits by cholinergic signaling can stabilize song behavior in adult male birds, which also provides a mechanism for the short-term plasticity and acute modulation of learned motor skills by cholinergic signaling in birds and even vertebrates. Further, we put forward a rational hypothesis concerning cholinergic modulation of song learning process.

## RESULTS

### Cholinergic signaling reduces RAPNs’ excitatory synaptic afferents in adult male and female zebra finches via mAChRs rather than nAChRs

To investigate the cholinergic modulation of RAPNs’ excitatory synaptic afferents in adult zebra finches, we first respectively recorded the influences of cholinergic receptor agonists and antagonists on RAPNs’ mEPSCs. The results showed that the non-selective cholinergic receptor agonist carbachol (CAR, 30 μM) significantly decreased the frequency and amplitude of RAPNs’ mEPSCs in both males (Figure 1B) and females (Figure 2B), indicating that CAR can reduce RAPNs’ excitatory synaptic afferents in adult male and female zebra finches.

**Figure 2.**
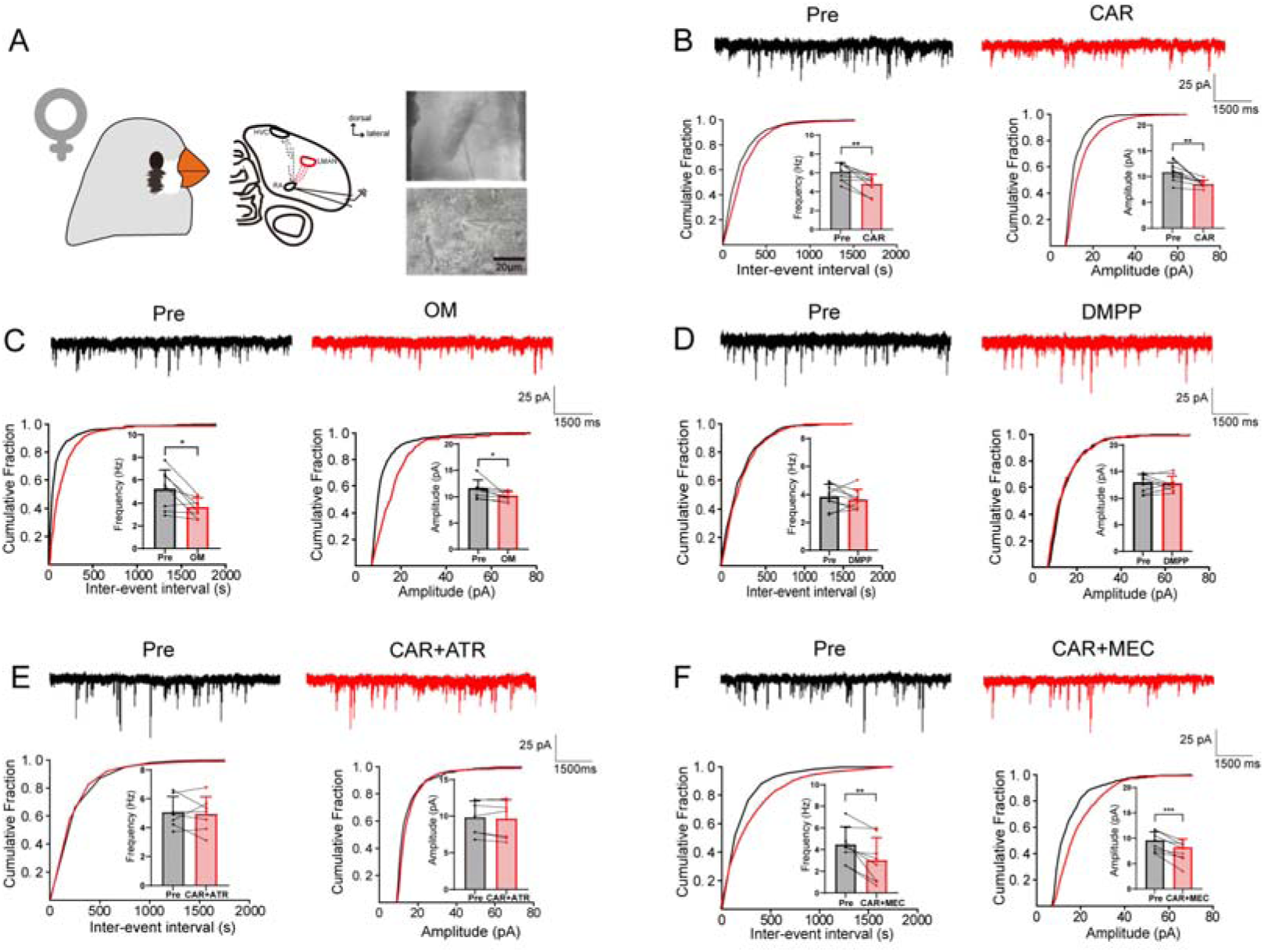
Cholinergic modulation of RAPNs’ mEPSCs in adult female zebra finches. (A) Experiment schematic. The pictures show in vitro whole-cell patch clamp recording of RAPNs’ mEPSCs in adult females. (B) Top, example traces of adult female RAPNs’ mEPSCs in Pre and CAR; Bottom, quantification of mEPSCs’ frequency (left) and amplitude (right) in Pre and CAR, n = 9. (C-F) Similar to (B). OM (C), n = 8; DMPP (D), n = 9; CAR+ATR (E), n = 7; CAR+MEC (F), n = 8. The variable n refers to the number of cells. * *p* ≤ 0.05, ** *p* ≤ 0.01, *** *p* ≤ 0.001, paired *t*-test.

The mAChR agonist Oxotremorine M iodide (OM, 10 μM) mimicked the effects of CAR, which also significantly reduced the frequency and amplitude of RAPNs’ mEPSCs in both males (Figure 1C) and females (Figure 2C). However, the nAChR agonist 1,1-Dimethyl-4-phenylpiperazinium iodide (DMPP, 10 μM) had no effect on the frequency and amplitude of RAPNs’ mEPSCs in both males (Figure 1D) and females (Figure 2D). When CAR and the mAChR antagonist atropine (ATR, 10 μM) were simultaneously added, the effect of CAR was blocked by ATR, and no significant changes were observed in the frequency and amplitude of RAPNs’ mEPSCs in both males (Figure 1E) and females (Figure 2E). When CAR and the nAChR antagonist mecamylamine (MEC, 10 μM) were co-administered, the effect of CAR was not blocked by MEC, and the frequency and amplitude of RAPNs’ mEPSCs remained significantly decreased in both males (Figure 1F) and females (Figure 2F). This finding suggests that cholinergic signaling reduces RAPNs’ excitatory synaptic currents in adult male and female zebra finches via mAChRs but not nAChRs.

### Cholinergic signaling reduces HVC-RAPN excitatory synaptic transmission in adult male zebra finches via mAChRs rather than nAChRs

HVC-RA pathway is a crucial excitatory synaptic afferent of RA. Our further results showed that CAR significantly reduced the amplitude of RAPNs’ evoked EPSCs (eEPSCs) induced by stimulating HVC-RA projection fibers in adult males (Figure 3B), indicating that CAR can prominently decrease HVC-RAPN excitatory synaptic transmission. Meanwhile, the effect of CAR was accompanied by a significant increase in paired-pulse facilitation ratio (PPR) (Figure 3B), suggesting that presynaptic mechanism was involved in the role of CAR on HVC-RAPN excitatory synaptic transmission.

**Figure 3.**
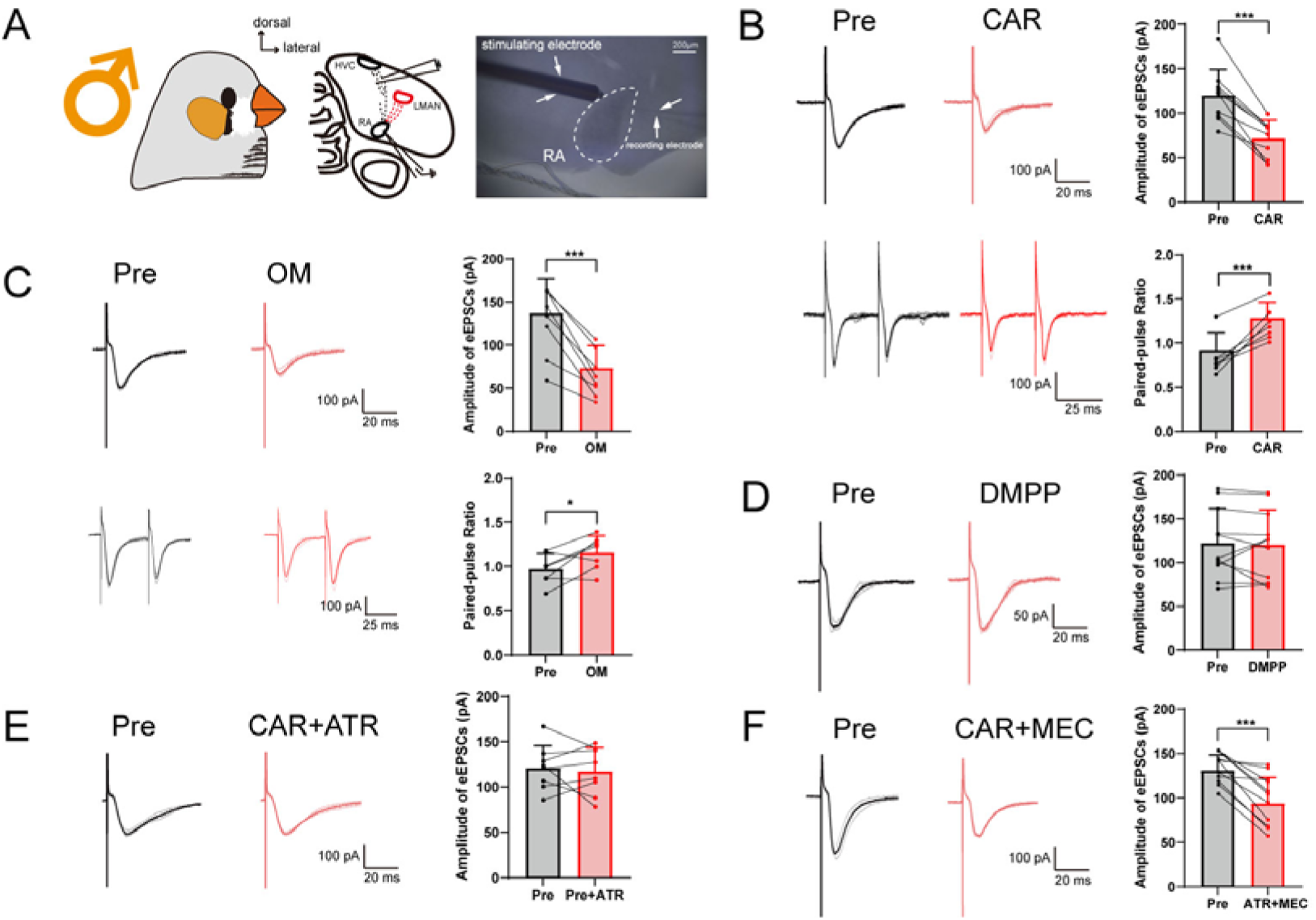
Cholinergic modulation of HVC-RAPN eEPSCs in adult male zebra finches. (A) Experiment schematic. The picture shows the location of the stimulating electrode in in vitro whole-cell patch clamp recording of HVC-RAPN eEPSCs in adult males. (B) Top, example traces of adult male HVC-RAPN eEPSCs in Pre and CAR. The transparent curves represent individual EPSCs, and the bold curves on top represent the average across trials. Quantification of eEPSCs’ amplitude in Pre and CAR, n = 10; Bottom, example traces of eEPSCs in paired pulse stimulation. PPR quantification in Pre and CAR, n = 7. (C) Similar to (B). OM: Top, n = 8; Bottom, n = 7. (D-F) Example traces of adult male HVC-RAPN eEPSCs in Pre, DMPP (D), CAR+ATR (E) and CAR+MEC (F). Quantification of eEPSCs’ amplitude in Pre, DMPP (D, n = 11), CAR+ATR (E, n = 8), and CAR+MEC (F, n = 11). The variable n refers to the number of cells. * *p* ≤ 0.05, *** *p* ≤ 0.001, paired *t*-test.

OM mimicked the effect of CAR, which likewise significantly reduced HVC-RAPN eEPSCs’ amplitude (Figure 3C) and caused a significant elevation in PPR (Figure 3C). However, DMPP had no effect on HVC-RAPN eEPSCs’ amplitude (Figure 3D). If CAR and ATR were concurrently administered, the inhibitory action of CAR on HVC-RAPN eEPSCs’ amplitude was obstructed by ATR (Figure 3E). Nevertheless, when CAR and MEC were concurrently administered, MEC failed to neutralize the effect of CAR (Figure 3F). These results suggest that cholinergic signaling reduces HVC-RAPN excitatory synaptic transmission in adult male zebra finches via mAChRs but not nAChRs.

### Cholinergic signaling reduces LMAN-RAPN excitatory synaptic transmission in adult male zebra finches via nAChRs rather than mAChRs

LMAN-RA pathway is another important excitatory synaptic afferent of RA. CAR significantly decreased the amplitude of RAPNs’ eEPSCs elicited by stimulating LMAN-RA projection fibers in adult males (Figure 4B), indicating that CAR can markedly reduce LMAN-RAPN excitatory synaptic transmission. During the CAR effect, PPR was significantly increased (Figure 4B), suggesting that presynaptic mechanism was involved in the role of CAR on LMAN-RAPN excitatory synaptic transmission.

**Figure 4.**
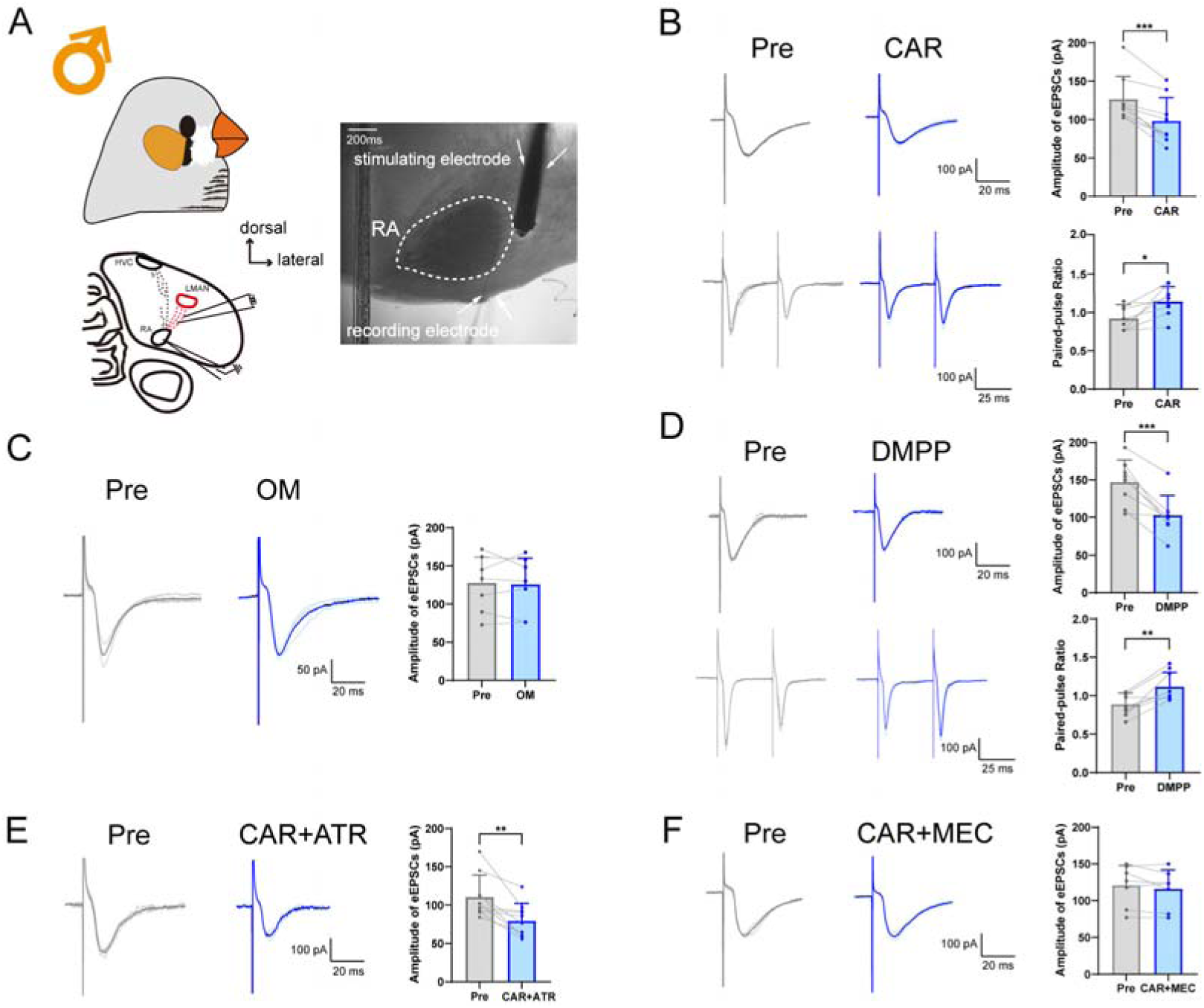
Cholinergic modulation of LMAN-RAPN eEPSCs in adult male zebra finches. (A) Experiment schematic. The picture shows the location of the stimulating electrode in in vitro whole-cell patch clamp recording of LMAN-RAPN eEPSCs in adult males. (B) Top, example traces of adult male LMAN-RAPN eEPSCs in Pre and CAR. The transparent curves represent individual EPSCs, and the bold curves on top represent the average across trials. Quantification of eEPSCs’ amplitude in Pre and CAR, n = 9; Bottom, example traces of eEPSCs in paired pulse stimulation. PPR quantification in Pre and CAR, n = 8. (C) Example traces of adult male LMAN-RAPN eEPSCs in Pre and OM. Quantification of eEPSCs’ amplitude in Pre and OM, n = 8. (D) Similar to (B). DMPP: Top, n = 8; Bottom, n = 8. (E-F) Similar to (C). CAR+ATR (E), n = 10; CAR+MEC (F), n = 8. The variable n refers to the number of cells. * *p* ≤ 0.05, ** *p* ≤ 0.01, *** *p* ≤ 0.001, paired *t*-test.

OM exerted no influence on LMAN-RAPN eEPSCs’ amplitude (Figure 4C). However, DMPP simulated the effect of CAR, significantly reducing LMAN-RAPN eEPSCs’ amplitude (Figure 4D), and simultaneously causing a significant increase in PPR (Figure 4D). When CAR and ATR were applied together, the reduction effect of CAR on LMAN-RAPN eEPSCs’ amplitude was not blocked by ATR (Figure 4E). If CAR and MEC were applied simultaneously, the effect of CAR was blocked by MEC (Figure 4F). These results demonstrate that cholinergic signaling reduces LMAN-RAPN excitatory synaptic transmission in adult male zebra finches via nAChRs but not mAChRs. Namely, receptor mechanisms of cholinergic modulation on the excitatory synaptic transmission of the two song premotor pathways, LMAN-RA and HVC-RA, in males are different.

### Cholinergic signaling has no effect on HVC-RAPN excitatory synaptic transmission in adult female zebra finches

In contrast to the findings in males, CAR exerted no impact on HVC-RAPN eEPSCs’ amplitude in adult females (Figure 5B). Additionally, neither DMPP (Figure 5C) nor OM (Figure 5D) had any effect on HVC-RAPN eEPSCs’ amplitude. This postulates that HVC-RAPN excitatory synaptic transmission in adult female zebra finches is not regulated by cholinergic signaling.

**Figure 5.**
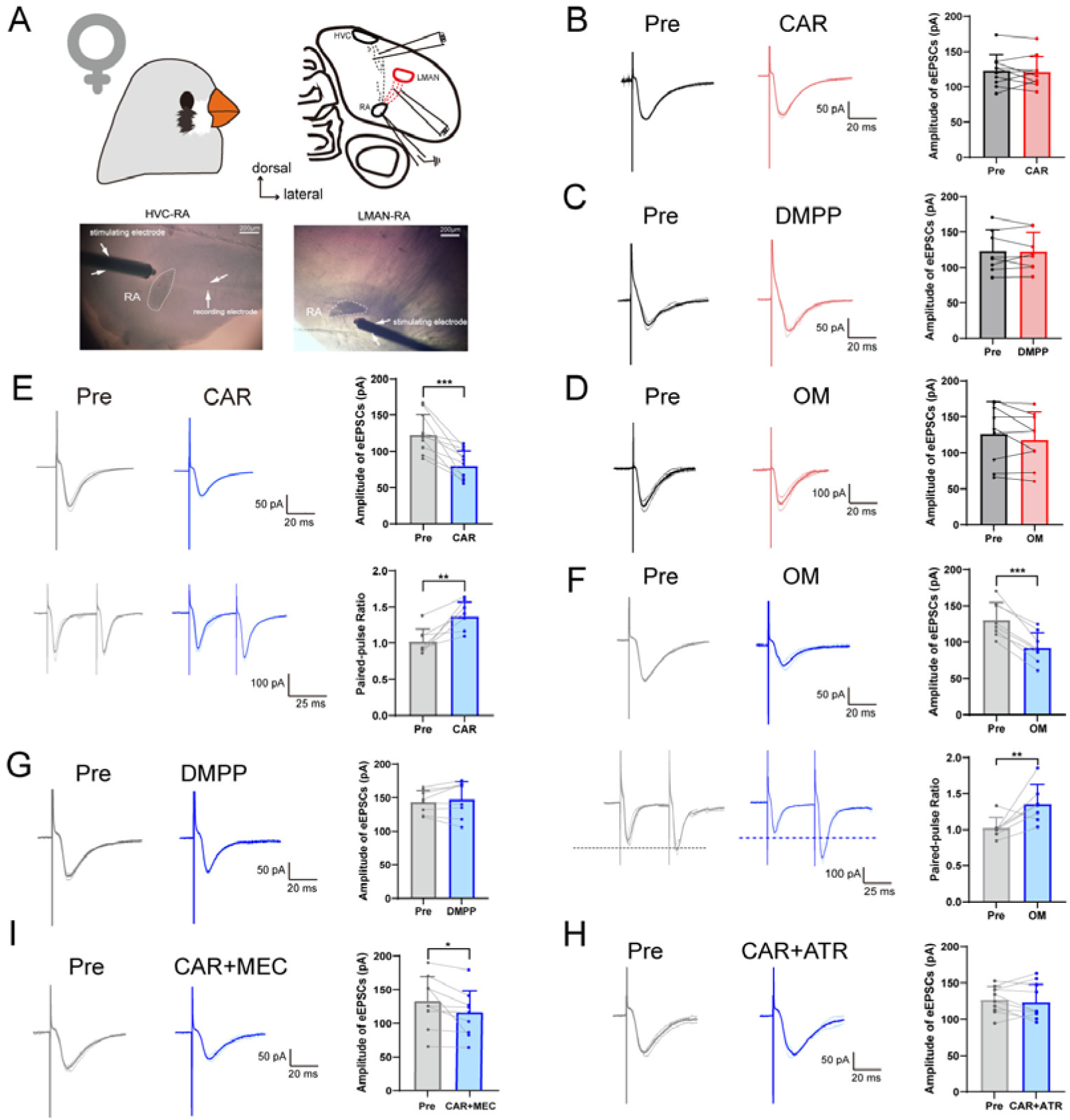
Cholinergic modulation of HVC-RAPN and LMAN-RAPN eEPSCs in adult female zebra finches. (A) Experiment schematic. The pictures respectively show the locations of the stimulating electrodes in in vitro whole-cell patch clamp recording of HVC-RAPN (left) and LMAN-RAPN (right) eEPSCs in adult females. (B) Example traces of adult female HVC-RAPN eEPSCs in Pre and CAR. The transparent curves represent individual EPSCs, and the bold curves on top represent the average across trials. Quantification of eEPSCs’ amplitude in Pre and CAR, n = 10. (C-D) Similar to (B). DMPP (C), n = 8; OM (D), n = 9. (E) Top, example traces of adult female LMAN-RAPN eEPSCs in Pre and CAR. Quantification of eEPSCs’ amplitude in Pre and CAR, n = 10; Bottom, example traces of eEPSCs in paired pulse stimulation. PPR quantification in Pre and CAR, n = 7. (F) Similar to (E). OM: Top, n = 8; Bottom, n = 7. (G) Example traces of adult female LMAN-RAPN eEPSCs in Pre and DMPP. Quantification of eEPSCs’ amplitude in Pre and DMPP, n = 8. (H-I) Similar to (G). CAR+ATR (H), n = 10; CAR+MEC (I), n = 9. The variable n refers to the number of cells. * *p* ≤ 0.05, ** *p* ≤ 0.01, *** *p* ≤ 0.001, paired *t*-test.

### Cholinergic signaling reduces LMAN-RAPN excitatory synaptic transmission in adult female zebra finches via mAChRs rather than nAChRs

Similar to males, CAR significantly reduced LMAN-RAPN eEPSCs’ amplitude in adult females (Figure 5E), and led to a significant increase in PPR (Figure 5E), suggesting that there are also presynaptic mechanisms involved in the CAR modulation of LMAN-RAPN excitatory synaptic transmission in females.

OM mimicked the reduction effect of CAR on LMAN-RAPN eEPSCs’ amplitude in females (Figure 5F), accompanied by a significant increase in PPR (Figure 5F). Whereas, DMPP had no influence on LMAN-RAPN eEPSCs’ amplitude in females (Figure 5G). Simultaneously applying CAR and ATR, the effect of CAR was blocked by ATR (Figure 5H). While simultaneously applying CAR and MEC, the effect of CAR was not blocked by MEC (Figure 5I). These results indicate that, contrary to the receptor mechanism of cholinergic modulation on LMAN-RAPN excitatory synaptic transmission in males, cholinergic signaling reduces LMAN-RAPN excitatory synaptic transmission in adult female zebra finches via mAChRs but not nAChRs.

### Cholinergic signaling within RA enhances song stability of adult male zebra finches via mAChRs rather than nAChRs

To further validate the impact of cholinergic signaling within RA on adult song behavior and its receptor mechanisms, the results of behavioral experiments combined with an in vivo pharmacological method (Figure 6A and C) showed that CAR (1 mM) microinjection onto RA significantly elevated the similarity to own song of adult males (Figure 6B and D) and reduced the song entropy (Figure 6B and E). OM (1 mM) microinjection onto RA exhibited a comparable effect to CAR, which also increased song similarity (Figure 6B and D) and decreased song entropy (Figure 6B and E), whereas DMPP (1 mM) microinjection had no effect on song similarity (Figure 6B and D) and entropy (Figure 6B and E). For comparison, the results of phosphate buffered saline (PBS) microinjection showed that there was no obvious change in song similarity (Figure 6B and D) and entropy (Figure 6B and E), thereby ruling out the influence brought by the microinjection operation per se. In addition, the sum of duration of all motif syllables, the sum of all gap durations and other song acoustic characteristics were not notably influenced by the diverse drug microinjections (Figure 6B, F, G, and Supplementary Figure 1). These results manifest that cholinergic signaling within RA enhances song stability of adult male zebra finches via mAChRs but not nAChRs. Moreover, short calls in males were not influenced by CAR within RA (Supplementary Figure 2).

**Figure 6.**
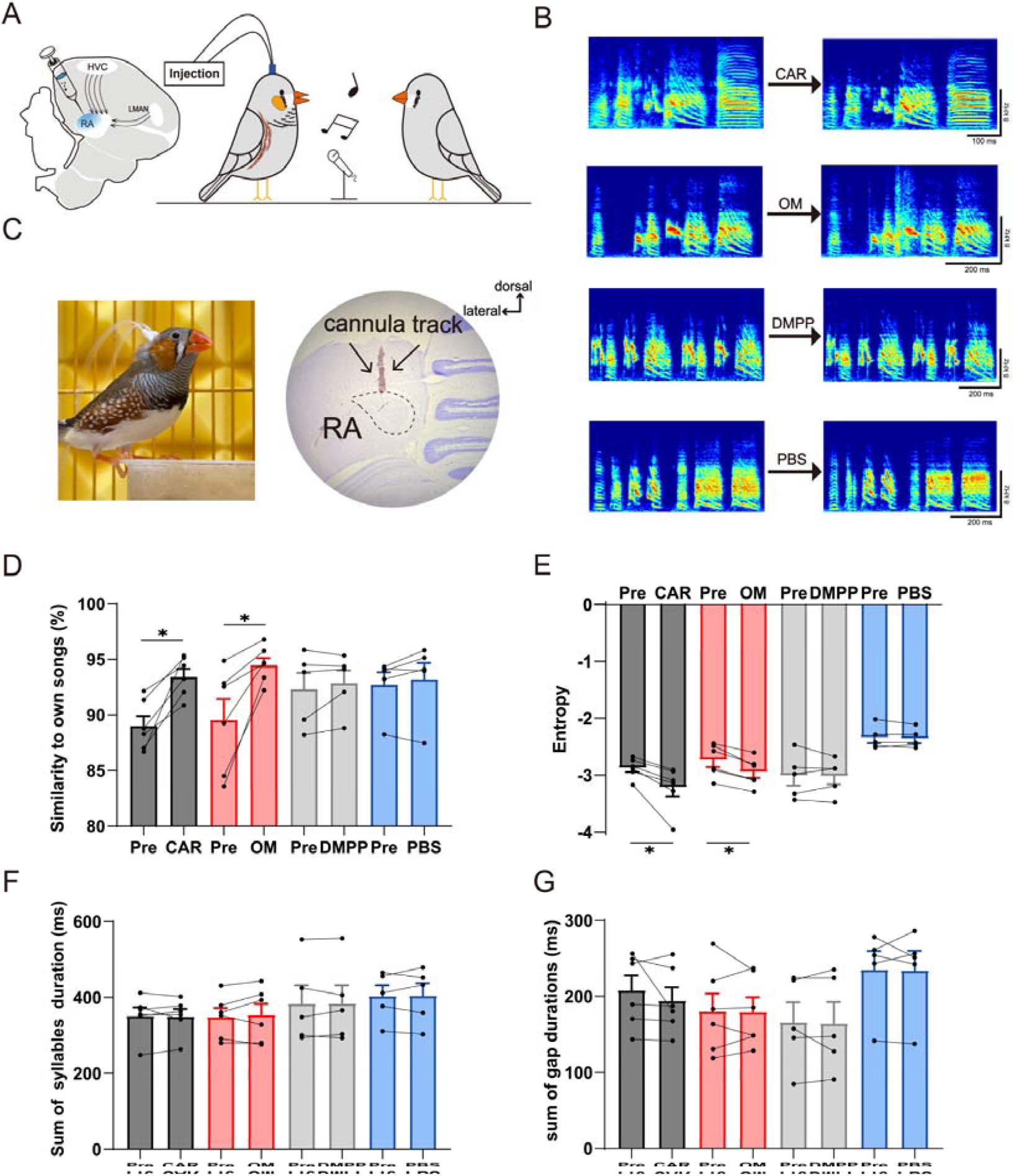
Cholinergic signaling within RA enhances song stability of adult male zebra finches via mAChRs rather than nAChRs. (A) Experiment schematic. In vivo pharmacological microinjection onto bilaterally RA and song recording. (B) Example spectrograms of motifs in Pre and CAR, Pre and OM, Pre and DMPP, Pre and PBS. (C) Left, a free-moving adult male zebra finch in a recording studio with cannulas bilaterally implanted over RA; Right, a Nissl-stained single-side coronal section containing RA and the cannula track. (D) Similarity to own songs in Pre and CAR (n = 6), Pre and OM (n = 6), Pre and DMPP (n = 5), Pre and PBS (n = 5). (E) Song entropy in Pre and CAR (n = 6), Pre and OM (n = 6), Pre and DMPP (n = 5), Pre and PBS (n = 5). (F-G) The sum of duration of all motif syllables and the sum of all gap durations were not significantly affected by CAR (n = 6), OM (n = 6), DMPP (n = 5), and PBS (n = 5). * *p* ≤ 0.05, Signed-rank test.

## DISCUSSION

To illuminate cholinergic modulation of songbirds’ song behavior, it is indispensable to elucidate the cholinergic regulatory mechanism at the neural pathway level. We discover a mechanism by which cholinergic signaling separately regulates song premotor circuits via different receptors to stabilize the songs of adult males. Although females do not sing, a similar mechanism exists in the same circuits of adult females. Furthermore, we postulate a hypothesis concerning cholinergic regulatory process of song learning in juveniles and song maintenance in adults. From a broader perspective, our findings provide significant insights into cholinergic regulation of learned motor skills.

### Songbirds are an excellent animal model for revealing cholinergic modulation of learned motor skills

Central cholinergic system exists in the brains of fish^23, 24, 25^, amphibians^26, 27^, reptiles^26, 27^, mammals^28, 29, 30, 31^, and birds^32, 33, 34, 35^, which mainly participates in various higher neural activities such as motor skill acquisition^3, 36^, learning and memory^37, 38^, sleep and wakefulness^39, 40^, and attention maintenance^41, 42^. ACh exerts different effects by binding to two different types of receptors, mAChRs (G-protein coupled receptors) and nAChRs (ligand-gated ion channel receptors). Both receptors are expressed in all layers of the vertebrate cerebral cortex^23, 31, 43^, with different expression patterns among different layers and cell types^44^.

The organization of avian subcortical cholinergic system is analogous to that of mammals. Research has confirmed that the cholinergic system of pigeon in basal forebrain, epithalamus, isthmus, and hindbrain closely resembles that of reptiles and mammals^32^. Further research indicates that the basal telencephalic cholinergic system in pigeons and budgerigars is anatomically very similar to the cholinergic basal nuclear system in mammals, particularly in terms of their projection patterns in cerebral cortex or pallium^35, 45^. Although the neurons in avian pallium are not hierarchically organized like those in mammalian cerebral cortex, cholinergic receptor distribution in avian brain also shows regional specificity^33, 34^.

In songbirds, both male and female zebra finches have cholinergic projection fibers in song nuclei and auditory nuclei^46, 47^. Studies have shown that HVC and RA are rich in acetylcholinesterase, suggesting a strong cholinergic dominance over these two nuclei^15, 16, 46, 47, 48^. The cholinergic innervation of HVC and RA originates from VP, which is analogous to the mammalian basal forebrain cholinergic system, specifically the nucleus basalis of Meynert^12^.

Evidence suggests that mAChRs are expressed within song nuclei^49, 50^, and nAChRs are also present in multiple song nuclei^51^. Asogwa et al. cloned four out of the five mammalian mAChR subunits (Chrm2-5) in male zebra finches. They found that, at each developmental stage, the expression of excitatory subunits (chrm3 and chrm5) consistently exhibits lower levels compared to inhibitory subunits (chrm2 and chrm4) within most song nuclei of male zebra finches, including RA^15^. Recently, all 15 types of nAChR subunits in male zebra finch brain were cloned. It was confirmed that most nAChR subunits (except for ChrnA1, A6, A9, and A10) are expressed in song related pathways during the critical period of song learning, while only 6 nAChR subunits (ChrnA2-5, A7, and B2) are expressed in adulthood^16^. This suggests that the expression of most nAChR subunits undergoes continuous changes throughout song formation process in various song nuclei, and the expression types of nAChR subunits in adulthood are much fewer than those during song learning.

Central cholinergic system plays a crucial role in the regulation of learned motor skills^52, 53, 54^. However, the precise mechanisms through which this system governs such learning remain elusive, and elucidating its role poses significant challenges. Songbird’s song learning is a complex form of sensory-motor learning, which is an extremely rare vocal learning behavior similar to human language learning^5^. Pharmacological and behavioral research showed that subcutaneous administration of mid (0.18mg/kg) and high (0.54mg/kg) dose of the nAChR agonist nicotine can elicit a significant increase in the song production and locomotor activity of adult male zebra finches^55^. Continuous nicotine administration for 7 days can alter the tempo and rhythm of adult crystallized songs, and this effect may persist for an extended period (over 2 months) after discontinuation of nicotine administration^56^. These findings indicate that central cholinergic system likely plays a significant role in the vocalization and song learning processes of songbirds. Therefore, songbirds could serve as an excellent research model for in-depth exploration of cholinergic regulation on complex motor control and learning mechanisms, including human speech.

### A mechanism by which cholinergic signaling separately regulates song premotor circuits of male songbirds through different receptors thereby promoting song stability

The electrophysiological activities of HVC and RA are associated with the neural encoding of song motor instructions. The motor control signals generated by HVC require processing through RA in order to ultimately produce appropriate song motor encoding for coordinated activation of syringeal and respiratory muscles^57, 58^. Shea and Margoliash found that cholinergic signaling attenuates the auditory responses of HVC and RA to bird’s own song in adult male zebra finches^59^. Furthermore, distinct HVC projection neurons (HVC-to-RA and HVC-to-basal ganglia projection neurons) exhibit differential responses to cholinergic signaling, and this diversity of response may facilitate the coordination of cholinergic regulation between HVC-RA motor command transmission and the reception of HVC motor signals by AFP^60^. Jaffe and Brainard found that dialyzing CAR into HVC increases HVC neuron activities, resulting in an elevation of song pitch, amplitude, tempo and stereotypy to levels approaching those of direct courtship songs in adult male Bengalese finches, which can be weakened by blocking mAChRs^61^.

An early study revealed a notable elevation in ACh levels within HVC, LMAN and RA of zebra finches during the critical period for song acquisition, followed by a gradual decline as the birds approach adulthood^14^. ACh transient increase within RA during critical period is associated with heightened acetyltransferase (ChAT) activity^62^. Additionally, biochemical research has shown that during the critical period of song learning in zebra finches, CAR significantly increases the phosphoinositide turnover within the synaptoneurosomes of RA neurons^63^. Through intracellular electrophysiological recordings in brain slices, Salgado-Commissariat et al. discovered that nicotine enhances RA neuron excitability in adult male zebra finches, without distinguishing between the types of RA neurons (projection neurons or interneurons). Furthermore, by activating different nAChR subtypes, they demonstrated that tetanic stimulation induces bidirectional synaptic plasticity (long-term potentiation and long-term depression) in LMAN-RA pathway^64^.

RAPNs are homologous to layer 5 pyramidal neurons in mammalian motor cortex, and their activities govern song production^65, 66^. Our previous results obtained from brain slice whole-cell current-clamp recordings indicate that cholinergic signaling can induce a hyperpolarization of membrane potential, an increase in afterhyperpolarization potential and membrane conductance, and a reduction in action potential firing in RAPNs of adult male zebra finches via mAChRs rather than nAChRs^22, 67^. This suggests that under physiological conditions, cholinergic signaling may influence RAPN activities by altering their intrinsic membrane properties, thereby regulating song behavior. In present study, the results of brain slice whole-cell voltage-clamp recordings showed that cholinergic signaling significantly reduces RAPNs’ mEPSCs in adult male zebra finches through mAChRs rather than nAChRs, indicating that RAPNs’ excitatory synaptic afferents in adulthood is regulated by cholinergic signaling, and mAChR mediated action remains predominant. Cholinergic regulation on RAPNs’ intrinsic membrane properties and excitatory synaptic afferents is predominantly mediated by the inhibitory effect of mAChRs to reduce excitability, which is consistent with the circumstance that the expression of inhibitory mAChR subunits maintains higher levels compared to excitatory mAChR subunits within RA at all development stages^15^.

In mammals, similar to our results, it has been reported that CAR reduces glutamatergic excitatory synaptic transmission of layer 5 pyramidal neurons in rat sensory cortex, and this effect is blocked by mAChR antagonists^68^. In fact, the impacts of cholinergic signaling on neurons in diverse cortical areas and different layers are not homogeneous. It has been disclosed through pharmacological methods that nAChRs and mAChRs jointly contribute to mediating the ACh’s action on the neurons in layer 6 of primary motor cortex. However, the neurons in layer 6 of combined medial prefrontal cortex display a stronger response to ACh, and the effect is mainly mediated by nAChRs^29^. By comparing the aforementioned studies on mammals with our experimental results, it can be noted that the cholinergic regulation of song premotor brain regions in songbirds exhibits both similarities to that of cerebral cortex in mammals and interspecific dissimilarities or its functional specificity.

It is worth noting that the pharmacological research on layer 6A pyramidal cell types of rat somatosensory cortex in recent years has revealed that glutamate synaptic transmission in the corticocortical pyramidal cells is inhibited via M4 mAChRs, but in the corticothalamic pyramidal cells, it is enhanced via α4β2 nAChRs. This study indicates a potential mechanism through which ACh might separately regulate diverse projection neurons in the same cortical region via both nAChRs and mAChRs, thus achieving the coordination among synaptic transmission functions^44^. However, within the song motor nucleus RA of songbirds, there is merely one type of projection neurons (projecting to brainstem motor nuclei). The integration of song premotor signals from HVC-RA and LMAN-RA pathways within RAPNs is the critical determinant in generating the encoded output for controlling song production^69, 70^. The more crucial mechanism underlying cholinergic modulation of song behavior might lie in how to achieve the discrete regulation of HVC-RA pathway and LMAN-RA pathway, concomitant with the reciprocal coordination and collaboration in the regulation between the signal transmissions of the two pathways. Our present results revealed that cholinergic signaling markedly decreases glutamatergic HVC-RAPN eEPSCs’ amplitude via mAChRs in adult male zebra finches. On the contrary, the reduction of glutamatergic LMAN-RAPN eEPSCs’ amplitude caused by cholinergic signaling was mediated by nAChRs in adult males. Meanwhile, cholinergic regulation of both HVC-RAPN and LMAN-RAPN synaptic transmission is accompanied by alterations in PPR (an electrophysiological indicator suggesting presynaptic mechanisms^71^). Our discoveries demonstrate a mechanism by which cholinergic signaling achieves the distinct regulation of song premotor circuits through two receptors, nAChRs and mAChRs. Recently, it has also been found in adult mice that cholinergic signaling can differentially regulate the glutamatergic synaptic transmission of the projections from prelimbic cortex and thalamus to pyramidal neurons of basolateral amygdala, respectively, through different receptors^72^. The disclosure of the differential regulatory mechanism of cholinergic receptors further underpins the crucial role of cholinergic regulation adapted to specific functions in relevant brain regions.

Most tellingly, our further behavioral research combined with in vivo targeted pharmacological manipulation indicated that CAR microinjection onto RA significantly enhances song stability of adult male zebra finches. The result is consistent with the finding of Jaffe and Brainard in adult male Bengalese finches that dialysis infusion of CAR into HVC affects singing, both increasing song stability^61^. Furthermore, microinjection of mAChRs agonist has the same effect as CAR, whereas microinjection of nAChRs agonist has no effect. Therefore, the result indicates that cholinergic signaling within RA governs song behavior in adulthood predominantly via mAChRs rather than nAChRs. This receptor mechanism is consistent with that of cholinergic signaling regulating RAPNs’ mEPSCs and intrinsic membrane properties in adult males, all through mAChRs but not nAChRs^22^. Taken together, given that RAPNs are characterized by the electrophysiological characteristic of high excitability^73, 74^, cholinergic signaling may appropriately lower the high excitability of RAPNs by diminishing the excitatory synaptic transmission of song premotor pathways in adult males via mAChRs, thereby improving signal-to-noise ratio to enhance their information integration capacity and ultimately promoting song stability. Moreover, evidence indicates that apart from cholinergic signaling, RA is also controlled by a multitude of other neurotransmitters, such as monoamine neurotransmitters^75, 76, 77, 78, 79^. Cholinergic signaling could merely be one constituent in the modulation of the chemical “cocktail”. Meanwhile, RAPNs’ inhibitory synaptic inputs from the GABAergic interneuron network would exert an important balancing effect^80^.

Another investigation has validated that cholinergic signaling within RA also has a substantial impact on song learning during developmental stage. Puzerey et al. employed in vivo reverse microdialysis technique to administer nAChR and mAChR antagonists in juvenile zebra finch RA for the purpose of chronically blocking cholinergic signaling in RA during song learning. After several weeks, it was discovered that singing quantity declined, and song learning process became disordered, such as excessive increase in song variability, abnormal acoustic features, and a decrease in similarity to tutor song^17^. It is currently known that LMAN-RA pathway assumes a pivotal role in song learning^81, 82^ and may provide motor correction signals for HVC-RA pathway^58, 83, 84^. The lesion of LMAN in juvenile zebra finches can disrupt song acquisition, while the lesion of LMAN in adulthood has no impact on the already stable songs^85, 86^. Furthermore, throughout the process of development and song acquisition, the synaptic density and amount of HVC-RA projection fibers increase conspicuously, yet the synaptic density and amount of LMAN-RA projection fibers diminish markedly^11, 87^. Consequently, based on the findings of predecessors and ours, it is reasonable to believe that during the song learning period of juvenile zebra finches, the mutual coordination and collaboration between the cholinergic regulation of LMAN-RA pathway mediated by nAChRs and that of HVC-RA pathway mediated by mAChRs exert a significantly facilitating effect on the process of song learning. However, during the process of song gradually stabilizing, with the weakening of LMAN-RA pathway and the strengthening of HVC-RA pathway, the cholinergic modulation of LMAN-RA pathway mediated by nAChRs also weakens accordingly. The finding of Asogwa et al. that the expression types of nAChR subunits in adulthood within song related pathways, including LMAN-RA pathway, are much fewer than those in critical period of song learning offers support for our result^16^. Therefore, the reinforcing effect of cholinergic signaling on the stability of adult birdsongs is mainly regulated by HVC-RA pathway mediated via mAChRs (Figure 7A).

**Figure 7.**
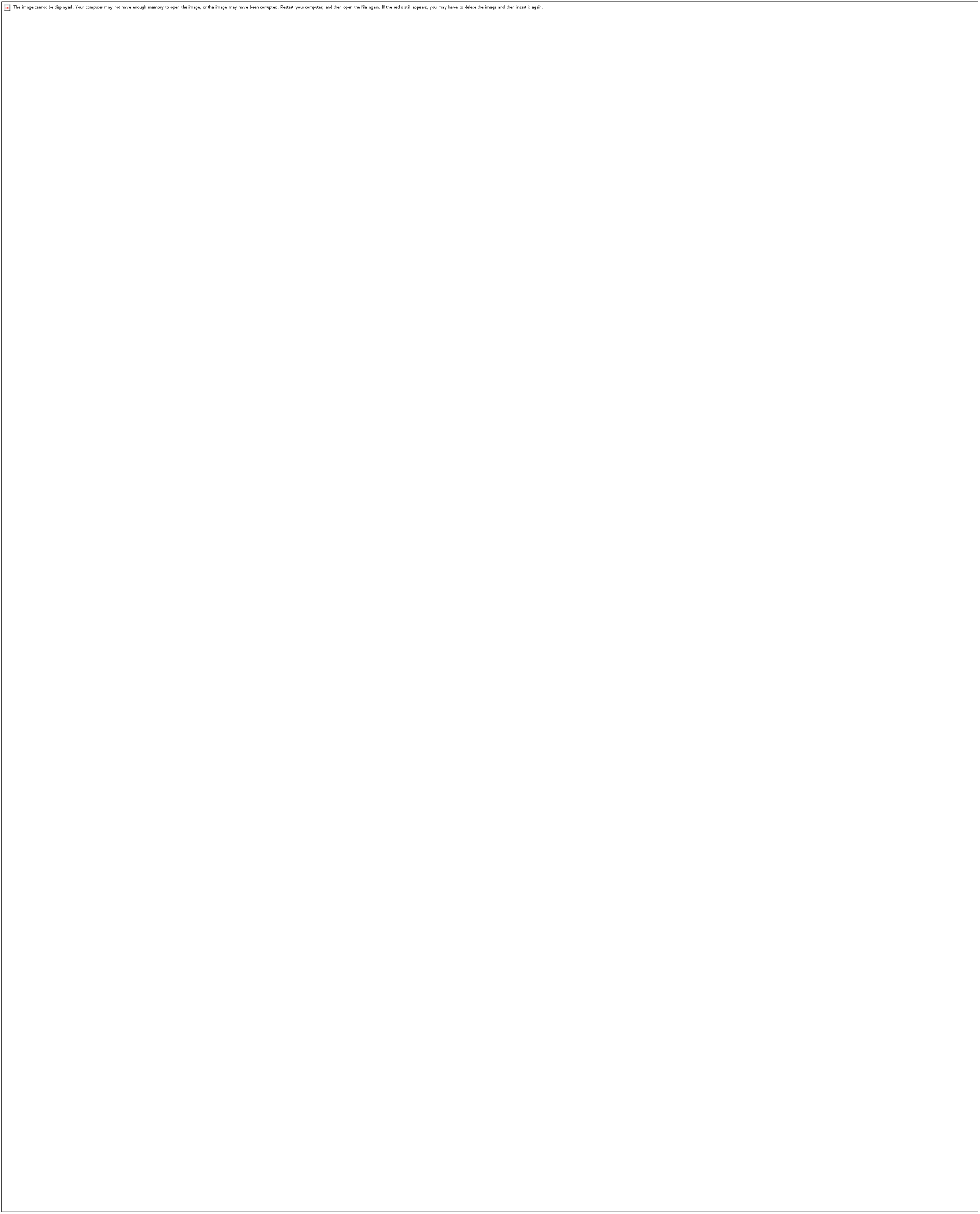
Mechanism hypotheses on the cholinergic modulation of song premotor circuits in adult male and female zebra finches. (A) A hypothesis regarding the mechanism that cholinergic signaling influences song stability by regulating song premotor circuits in adult males. (B) A hypothesis regarding the mechanism that cholinergic signaling impacts song perception and/or selection of adult females. Details in “DISCUSSION”.

Although cholinergic modulation of LMAN-RA pathway mediated by nAChRs turns quiescent or latent in adulthood, our experimental observations still imply that in the context of electrical stimulation of LMAN-RA projection fibers, cholinergic signaling can give rise to a decline in LMAN-RAPN excitatory synaptic transmission in adult males via nAChRs. It is suggested that this pathway may undergo adaptive changes under the elicitation of certain physiological conditions (such as experience-dependent song plasticity^88^) or pathological conditions. For example, in the event that the auditory feedback of adult songbirds undergoes modifications or is absent, LMAN-RA pathway will facilitate variations in the signal output of HVC-RA pathway and induce changes in vocalization^89^. It was reported that by activating different nAChR subtypes, long-term synaptic plasticity could be induced in the LMAN-RA pathway of adult male zebra finches through tetanic stimulus^64^, suggesting the potential cholinergic regulation of LMAN-RA pathway in adulthood.

### Cholinergic modulation of LMAN-RA pathway mediated by mAChRs may be a potential mechanism for perception and selection of birdsongs in females

Zebra finches exhibit distinct sexual dimorphism in both appearance and song behavior. Adult male birds attract females through singing, whereas females do not sing^90^. Correspondingly, most of song control nuclei in males are much larger than those in females^91^. Studies have demonstrated that the neurons within HVC and RA of females are smaller and less than those of males, so the volumes of the two nuclei in females are considerably smaller than those in males^92^. Such a discrepancy is associated with sex hormones^93^, and may even be linked to sex-specific transcriptomes^8^. It has been reported that RAPNs’ excitability of adult female zebra finches is significantly lower than that of males^94^, and there are sex differences in the intrinsic excitability of maturing RAPNs^95^. Our previous studies have demonstrated that in adult female zebra finches, both the frequency and amplitude of RAPNs’ spontaneous EPSCs (sEPSCs) and mEPSCs are notably lower than those of males^96^, and both the frequency and amplitude of spontaneous inhibitory postsynaptic currents (sIPSCs) and mIPSCs of RAPNs in females are also conspicuously lower than those in males^80^. A study has shown that neuropeptides expression in the neurons of forebrain song control nuclei in female zebra finches is significantly lower than that in males^97^. However, the in-depth research regarding the effect of neurotransmitters on the song related nuclei of female songbirds is rather limited.

Previous immunohistochemical results demonstrated that the size and density of ChAT-immunoreactive somata projected from VP to HVC and RA in males are significantly higher than those in females^98^, indicating sex differences in the cholinergic modulation of song control system. Our present study showed that, similar to males, cholinergic signaling can significantly reduce RAPNs’ mEPSCs in adult female zebra finches via mAChRs rather than nAChRs, indicating that RAPNs’ excitatory synaptic afferents in adult females is regulated by cholinergic signaling, and the role of mAChRs predominates. To our knowledge, this should be the first time to provide direct evidence for cholinergic modulation of female song control nuclei.

Studies have indicated that the development and formation of HVC-RA pathway in female zebra finches is incomplete^99, 100, 101^. It has been reported that sex differences in the myelination of zebra finch HVC-RA tract are significant, and the degree of myelination in females is lower than that in males and remains unchanged with development^102^. Moreover, the quantity of RAPNs in females is markedly less than that in males^103^. These pieces of evidence seemingly offer a rational explanation for the fact that females do not sing. Nevertheless, it has still been proved that female zebra finches have HVC-RA projection fibers like males^104^. A study of Wang et al. demonstrated that electrical stimulation of HVC-RA projection fibers in adult female zebra finches could result in the corresponding neural activity in RA, indicating the presence of HVC-RA synaptic transmission in females^105^. In accordance with this finding, our present results indicate that RAPNs’ eEPSCs can be recorded by electrically stimulating HVC-RA projection fibers in adult female zebra finches, reconfirming the presence of HVC-RA synaptic transmission in adult females. However, we found that, in contrast to males, cholinergic signaling has no effect on HVC-RAPN eEPSCs in females, suggesting that HVC-RAPN excitatory synaptic transmission of females is not regulated by cholinergic signaling. This result indicates that cholinergic modulation of HVC-RA pathway in adult zebra finches exhibits a typical sex dimorphism.

Among all song nuclei, only the LMAN of females is approximately the same in volume as that of males^106^. Consistent with males, the LMAN of female zebra finches likewise forms neural connections with RA in early developmental stage^100^. Moreover, a study has found that in adulthood, the number of synapses in the LMAN of females is significantly higher than that of males^107^. However, it has also been found that LMAN neurons projecting to RA in adult female zebra finches are much less than that in adult males, which might be caused by the massive loss of LMAN-to-RA projection neurons in females during the process of sexual differentiation and song learning^108^. Our further examination revealed that cholinergic signaling significantly reduces the amplitude of RAPNs’ eEPSCs evoked by electrically stimulating the LMAN-RA projection fibers of adult female zebra finches. This process is accompanied by the alteration of PPR, indicating that the presynaptic mechanism participated in cholinergic regulation on LMAN-RAPN excitatory synaptic transmission in females. However, unlike the cholinergic regulation of LMAN-RAPN synaptic transmission in males mediated by nAChRs, that in females is mediated by mAChRs rather than nAChRs, which is in line with the receptor mechanism of cholinergic regulation on RAPNs’ mEPSCs in females. Like zebra finches, in another songbird cowbirds (Molothrus ater), only males sing. Although females do not sing, they can distinguish males by perceiving birdsongs. Research has shown that the volume of female LMAN and the quantity of neurons therein are positively correlated with mating choices, indicating that non-singing females may perceive and choose male songs by LMAN^109^. Hence, female zebra finches, comparable to their male counterparts, might establish memories of song templates (songs heard during juvenile period, such as father’s song^110, 111^) in the early life stages and gradually develop song perception, among which LMAN may assume a pivotal role^107^. Based on this clue, the cholinergic modulation of LMAN-RA pathway in females might be implicated in the procedure of their perception and selection for male songs. Finally, we put forward a mechanism hypothesis that cholinergic signaling impacts song perception and/or selection of adult female zebra finches (Figure 7B).

### Limitations of the study

While the roles of different cholinergic receptors were unambiguously identified by pharmacological means, the impact of inhibitory interneuron network within RA on RAPNs could not be precluded in in vivo experiments. The cholinergic modulation of RAPNs’ inhibitory synaptic afferents and whether it enhances or attenuates the effect of cholinergic signaling on birdsongs require further investigation. Our hypothesis regarding the cholinergic receptor mechanism in the song learning process of juveniles also remains to be verified. Additionally, the hypothesis concerning the significance of cholinergic modulation on song premotor circuits in females still requires elaborately designed behavioral experiments for validation.

## Methods

### Ethics statement

All the experiments described in this study were approved by the Institutional Animal Care and Use Committee of Jiangxi Science and Technology Normal University (3601020137931).

### Animals

The adult male and female zebra finches (Taeniopygia guttata) used in the research were obtained from a reputable supplier. These birds were all over 120 days old and had been raised adaptively in a spacious aviary under the light/dark cycle of 14 hours of light and 10 hours of darkness at a temperature of 24[. The experimental animals were randomly assigned to experimental groups according to sex, and the sample processing order and positions did not influence experimental outcome. Valid data were randomly collected from a total of 90 adult males and 65 females. The sample size was selected based on the 3R (Reduction, Replacement and Refinement) principle of animal experiments, statistical criteria, and published works.

### Slice preparation

Following anesthesia by isoflurane inhalation (1.5%-2%), the brain was carefully extracted. Subsequently, the fresh brain in the ice-water mixed slice solution at pH 7.3-7.4 and osmolarity 330-340 mOsm, containing a saturated mixture of 95% O_2_ and 5% CO_2_, was sectioned coronally using a vibrating microtome (7000 smz; Campden Instruments, UK) at a thickness of 250 µm. The slice solution was composed of 62.5 mM NaCl, 5 mM KCl, 28 mM NaHCO_3_, 248 mM sucrose, 1.3 mM MgSO_4_·7H_2_O, 10 mM glucose, and 1.26 mM NaH_2_PO_4_·H_2_O. The slices were transferred to a holding chamber containing oxygenated artificial cerebrospinal fluid (ACSF) at 35°C. The ACSF was composed of 125 mM NaCl, 2.5 mM KCl, 1.2 mM MgSO_4_[7H_2_O, 1.27 mM NaH_2_PO_4_[H_2_O, 25 mM NaHCO_3_, 25 mM glucose, and 2.0 mM CaCl_2_, pH 7.3-7.4. The slices were incubated at least 0.5 h and equilibrated to room temperature prior to electrophysiological recording.

### Patch-clamp electrophysiology

The visualization of RA in brain slices was achieved using infrared differential interference contrast microscopy (IR-DIC) through a microscope (BX51WIF, Olympus, Tokyo, Japan). Whole-cell patch clamp recordings were conducted at a temperature of approximately 24°C. Slices were perfused with carbogen-bubbled ACSF at a rate of 1-2 ml/min, while neurons were visualized under a 40× water immersion lens. Patch pipettes were fabricated from standard borosilicate capillary glass (BF150-117-10, Sutter Instruments, CA, USA) using a Flaming/Brown micropipette puller (P-1000, Sutter Instruments, CA, USA). All recording pipettes had an open-tip resistance ranging from 3.0 to 6.0 MΩ in the bath solution. The patch pipettes were filled with a solution containing the following concentrations: 120 mM KMeSO_4_, 5 mM NaCl, 10 mM HEPES, 2 mM EGTA, 5 mM QX-314, 2 mM ATP, 0.3 mM GTP, and 0.1% biocytin (pH adjusted to 7.3-7.4; osmolarity set at 340 mOsm). The identification of RAPNs was based on the following points: (1) spontaneous activity in tight-seal mode (spontaneous firing rates of RAPNs: 7-10 Hz for males, 5-10 Hz for females; Supplementary Figure 3B and G); (2) intracellular staining of biocytin for morphological identification after the completion of electrophysiological recording (Supplementary Figure 3C and H) ^73, 112^. The images were obtained by a confocal microscope (STELLARIS 5 WELL, Leica, Wetzlar, Germany). The same methods were applied to both of males and females. However, compared with males, female RAPNs had less spontaneous activity and smaller waveforms, and their recordings were not as stable as those of males. Therefore, in order to improve the success rate of female RAPNs recording, we tried to make pipette resistances at 5.0-6.0 MΩ.

To record the miniature excitatory postsynaptic currents (mEPSCs) of RAPNs, 1 µM TTX (Tetrodotoxin, Sigma-aldrich, USA) was applied to the bath solution in order to block spontaneous events driven by intrinsic Na^+^ channel-mediated action potentials. During eEPSCs recordings of RAPNs, 150 µM PTX (Picrotoxin, Sigma-aldrich, USA) was applied to the bath solution to block GABAA receptor-mediated inhibitory synaptic currents. Whole-cell voltage-clamp recordings were implemented in accordance with standard operations. The holding potential was set at –75 or –80 mV. Pipette capacitance and series resistance were compensated, and the liquid junction potential was corrected online before recording. During the recording process, real-time monitoring of series resistance was carried out at intervals of 2 min. Recordings with a change in the series resistance greater than 25% were abandoned. The signals underwent sampling at 10 kHz and filtering at 2 kHz using the MultiClamp 700B amplifier (Molecular Devices, CA, USA). The data were acquired using Clampfit 10.7 (Molecular Devices, CA, USA) through a Digidata 1550B (Molecular Devices, CA, USA). 90 s were selected for analysis in each mEPSCs recording, which included 100-500 events. Analysis of mEPSCs was performed using MiniAnalysis 6.0 software (Synaptosoft, GA, USA). We set 5 times the root mean square of background noise as the threshold. Afterwards, events with clear baseline, obvious peak and smooth rise were reconfirmed by eyes before being included in the analysis. Inter-event interval (IEI) and amplitude for events in the control and drug-administered groups were compared. The significance levels of IEI and amplitude shifts were determined by employing the nonparametric Kolmogorov-Smirnoff (K-S) test to assess cumulative probability distributions.

EPSCs were evoked through a concentric needle electrode placed in the efferent projections from HVC (as shown in Figure 3A and 5A) or LMAN (as shown in Figure 4A and 5A)^113^. The anatomy of HVC-RA and LMAN-RA projections in males is based on the existing literature^114^. In Supplementary Figure 4A, an image of the coronal brain section of an adult female zebra finch displays female RA outline and the distribution of the fibers projecting around female RA that is extremely similar to that of males. The duration of the stimulation pulses is 0.1 ms, and their voltage range is 10-600 mV. Detecting stimulation intensity adapted to evoking appropriate amplitudes of EPSCs was determined as soon as possible through a test interval of 10-15 s. Compared with the previous study^115^, and based on our test on the stability of evoked EPSCs (Supplementary Figure 5), during the data collection process, each group was given three to six detecting stimulations before and after drug administration respectively, with an interval of 30 s between each stimulation. The paired-pulse ratio (PPR), which is the ratio of the amplitude of the second evoked EPSC to that of the first EPSC, was assessed at an inter-pulse interval of 50 ms. The stimulation intensity of each pulse in the paired pulses was the same as that of single detecting stimulation, and the repetition frequency is also the same. Alterations in PPR reflect involvement of presynaptic mechanisms. Although the afferents of RA from HVC and LMAN are anatomically distinguishable, we still further verified these two glutamatergic pathways of both males and females using pharmacological methods. The situation of males we verified is consistent with previous reports^113^. HVC-RAPNs EPSCs of males were weakly blocked by 25 µM NMDA receptor antagonist 2-amino-5-phosphonopentanoic acid (AP5), while LMAN-RAPNs EPSCs of males were mainly blocked by AP5 (Supplementary Figure 3A). The situation we demonstrated among the females indicates that the glutamatergic pharmacology of putative female HVC-RA and LMAN-RA projections is the same as that in males (Supplementary Figure 3F). Additionally, we also provided an example of recordings of a neuron outside of female RA showing no effect of stimulation on LMAN-RA fibers (Supplementary Figure 4B).

### Drug Application

CAR (non-selective cholinergic receptor agonist, 30 µM, MCE, NJ, USA); OM (an agonist of mAChRs, 10 µM, MCE, NJ, USA); ATR (an antagonist of mAChRs, 10 µM, MCE, NJ, USA); DMPP (an agonist of nAChRs, 10 µM, MCE, NJ, USA); MEC (an antagonist of nAChRs, 10 µM, MCE, NJ, USA). The effects of these drugs on RAPNs were assessed through bath perfusion.

### In vivo microinjection

Birds were anesthetized using isoflurane inhalation (1.5%-2%) and placed in a stereotaxic apparatus. Cannulas (length: 3.5 mm, outer diameter: 0.41 mm, RWD, Guangdong, China) were implanted above bilateral RA, and as close as possible to ensure that the microinjection drug could penetrate into RA (25 birds; head angle: 60°; M/L: ± 2.47 mm; A/P: 0.7 mm; deep: left 2.8 mm, right 3 mm). After birds recovered from surgery, micro injectors (length: 4.0 mm, outer diameter: 0.21 mm, RWD, Guangdong, China) were inserted into the cannulas and connected to a syringe pump (R462, RWD, Guangdong, China) through flexible tubing. To avoid damaging HVC by the cannula and to minimize the damage to the HVC-RA projections, the stereotaxic coordinates were determined through pre-experiments. The verification data indicate that the surgery did not have a significant impact on the song self-similarity (Supplementary Figure 6A). In vivo pharmacological experiments were conducted to test the effects of ACh receptor agonists on song production in four male groups. Microinjection of 1 mM CAR, 1 mM OM, 1 mM DMPP, and PBS (for control experiments) was performed in RA of the four groups respectively. Before drug administration, a baseline period of singing for 2-3 hours was recorded in the morning. Then, each group was microinjected with the corresponding test drug at a constant flow rate of 1-1.5 µL/ min, and song recording continued for 3-4 hours. After the experiment, the position of implanted cannulas was anatomically verified (Figure 6C). The data of 3 birds with misplaced cannulas were excluded, but were used to provide controls for the surgery and microinjection (Supplementary Figure 6B). Furthermore, we used a fluorescent substance Alexa Fluor™ 488 (10000 MW, 1 µM, Thermofisher, MA, USA) to simulate drug microinjection to verify that it can definitely penetrate into RA (Supplementary Figure 6C).

### Song recording

Song recordings were conducted in a recording studio with dimensions of 2.1 × 1.2 × 1 m^3^, equipped with a TAKSTAR directional microphone (Guangdong Victory Electronics Co. Ltd., Guangzhou, China; frequency range: 50-20000 Hz) and a glass window (20 × 40 cm^2^). During the recording sessions, a male bird was placed in a cage near the window within the recording studio and able to see a female bird positioned outside. Song capturing was performed using Cool Edit Pro 2.0 (sampling rate: 44100 Hz; channels: stereo; resolution: 16-bit). In the song recording of each bird, 30 or 60 typical motifs were collected respectively before and after drug administration to assess drug effect on song production. One bird had less than 30 motifs after the first CAR administration. After its song self-similarity returned to the baseline level on the second day (Supplementary Figure 6D), the drug was administrated again and motifs were re-collected. In addition, we also recorded some short calls along with song recording. Short calls with low fundamental frequency occur when birds are in close proximity with conspecifics^116^. In the CAR group, we identified separately 10 short calls in the recordings of each of the three birds before and after drug administration (i.e. a total of 30 calls in Pre and 30 calls in CAR), and analyzed drug effect on short calls of males.

### Analysis of song features

The Sound Analysis Pro 2011 (SAP 2011) and MATLAB (R2024b) were used for song analysis. The songs were preprocessed, including: (1) quantile normalization (normalizing the amplitude differences caused by the varying distances between the sound source and the microphone); (2) noise suppression and signal enhancement; (3) syllable segmentation. The self-similarity, entropy, pitch, duration of motifs, amplitude and frequency modulation (FM) were analyzed. For each bird, 30 or 60 typical motifs were chosen and calculated song self-similarities by averaging the song similarity scores of all the possible combinations before and after drug administration, respectively. Entropy is a measure of the width and uniformity of the power spectrum. In acoustic analysis, entropy can reflect the spectral distribution of the sound signal, and lower entropy signifies higher tonality^117^. Pitch is the fundamental frequency, reflecting the melodic structure and recognizability of a song^61^. The duration of a motif is measured from the beginning of the first syllable to the end of the last syllable in the sequence, including syllable duration and gap duration. The syllable duration represents the sum of the durations of all the syllables in a motif, and the same is for the gap duration. Amplitude refers to the relative intensity of sound, which is measured in dB scale. FM is estimated based on time and frequency derivatives at different frequencies, which is the estimated value of the (absolute) slope of frequency traces relative to the horizontal line.

### Quantification and statistical analysis

The operator performing in vitro patch-clamp recording, in vivo microinjection, song recording, and data analysis was unaware of the design of pharmacological experiments. Test drugs were prepared and supplied by another operator. Paired t-tests were conducted to statistically compare the data of each group before and after drug administration, while the data of song parameters were analyzed statistically using the Signed-rank test^61^. The software tools employed for the creation of experimental diagrams were Adobe Illustrator 2021 (Adobe, CA, USA), Origin Pro 8.0 (OriginLab, MA, USA), and Graphpad Prism 9.5 (GraphPad Software, CA, USA).

### Data availability

The original data generated during the current study are available from the lead contact upon requests. Any additional information required to reanalyze the data reported in this work is available from the lead contact upon requests.

### Code availability

This paper does not report original code.

## Acknowledgements

This research was funded by the National Natural Science Foundation of China (32160123), Jiangxi Provincial Key Project of Natural Science Foundation (20212ACB205002), and Jiangxi Provincial Key Laboratory of Organic Functional Molecules (2024SSY05141).

We would like to thank Dr. Dan Yuan at the Analytical & Testing Center of Jiangxi Science and Technology Normal University for the help on neuronal visualization analysis.

## Author contributions

Investigation, Ning Xu, Yutao Zhang, and Yalun Sun; methodology, Wei Meng; data curation, Ning Xu, Yutao Zhang, Yalun Sun, and Xueqing Song; formal analysis, Wei Meng, Ning Xu, YangYang Cao, Xinqi Yang, and Yuejun Zhou; writing – original draft, Wei Meng and Ning Xu; writing – review & editing, Wei Meng and Ning Xu; conceptualization, resources, supervision, project administration, and funding acquisition, Wei Meng.

## Supplementary information

Supplementary Figure 1. Other song acoustic characteristics are not influenced by cholinergic signaling within RA.

Supplementary Figure 2. Short calls of males are not influenced by cholinergic signaling within RA.

Supplementary Figure 3. Pharmacological validation of HVC-RA and LMAN-RA pathways, as well as electrophysiological and morphological identification of RAPNs and RA interneurons.

Supplementary Figure 4. Female HVC-RA and LMAN-RA projections, and recordings of female RA peripheral neurons stimulated on LMAN-RA fibers.

Supplementary Figure 5. Examples of the stability test for EPSCs recordings show the reliable amplitudes of evoked EPSCs in both males and females.

Supplementary Figure 6. Validation of surgery impact, misplaced cannulas, drug diffusion and baseline before the 2nd administration.

## Lead contact

Further information and requests for data should be directed to and will be fulfilled by the lead contact, Wei Meng.

**Supplementary Figure 1.**
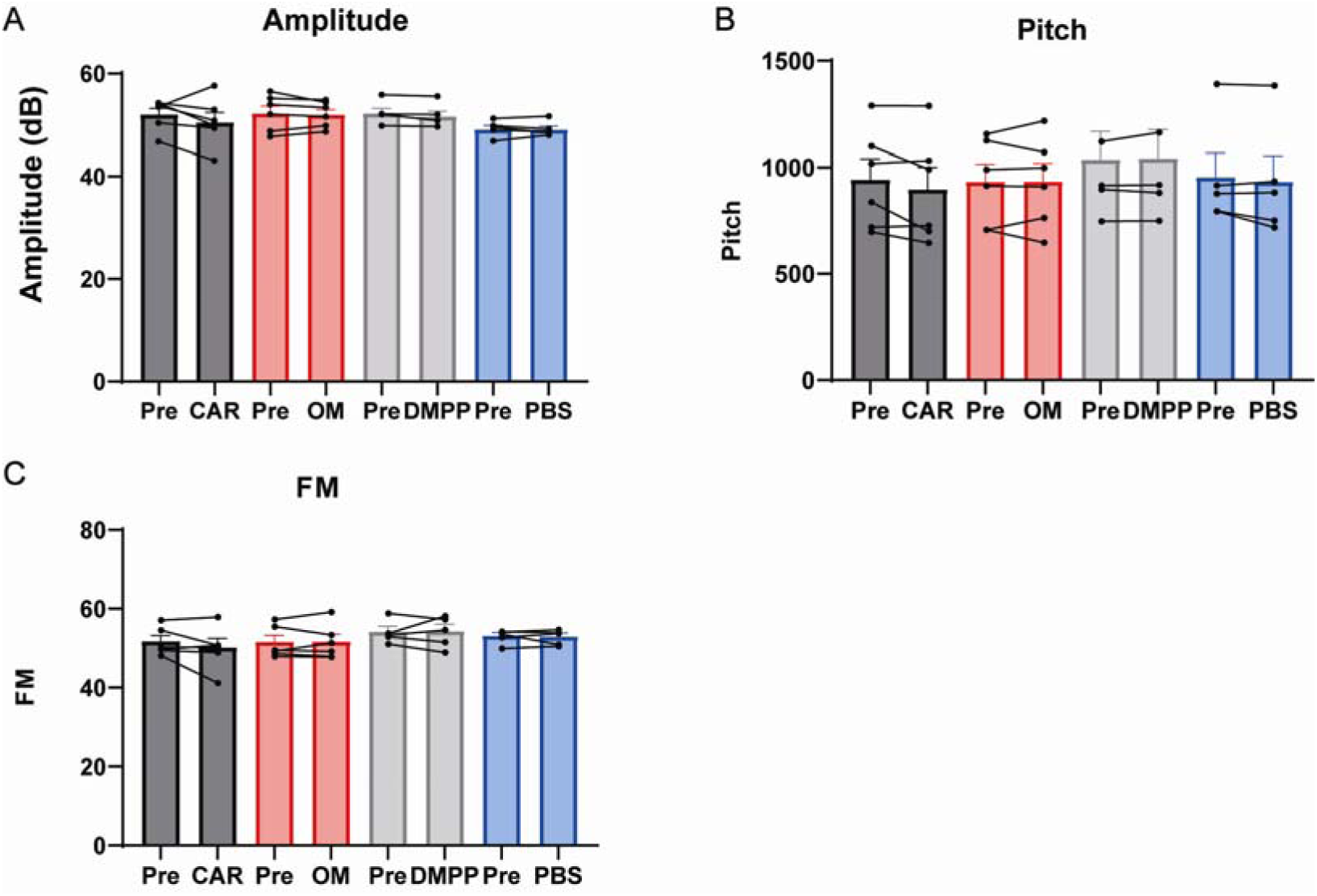
Other song acoustic characteristics are not influenced by cholinergic signaling within RA. **(A).** Amplitude in Pre and CAR (n = 6), Pre and OM (n = 6), Pre and DMPP (n = 5), Pre and PBS (n = 5) **(B).** Pitch in Pre and CAR (n = 6), Pre and OM (n = 6), Pre and DMPP (n = 5), Pre and PBS (n = 5). **(C).** FM in Pre and CAR (n = 6), Pre and OM (n = 6), Pre and DMPP (n = 5), Pre and PBS (n = 5). Signed-rank test.

**Supplementary Figure 2.**
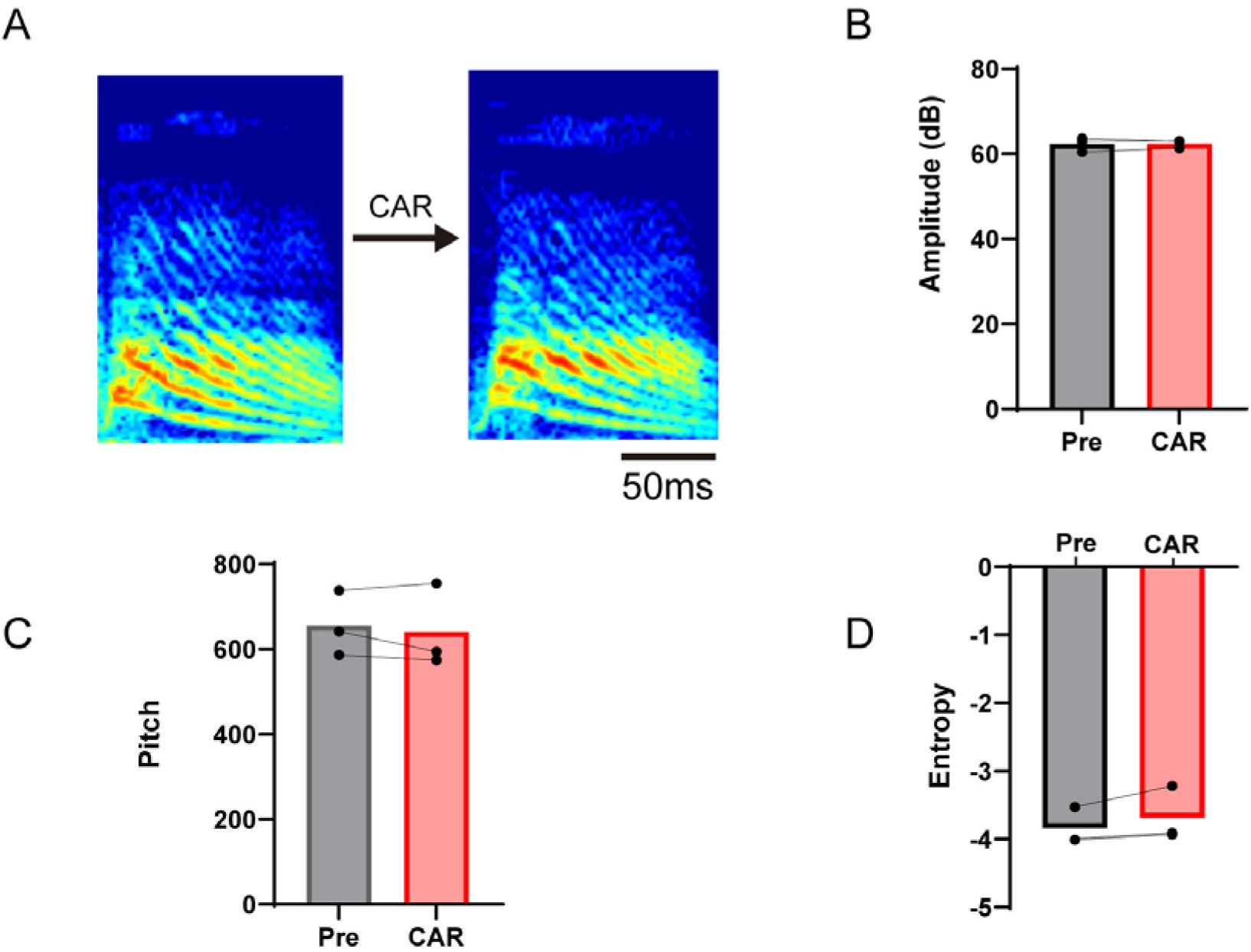
Short calls of males are not influenced by cholinergic signaling within RA. **(A).** Example spectrograms of male short calls in Pre and CAR. **(B).** Amplitude of male short calls in Pre and CAR (n = 3). **(C).** Pitch of male short calls in Pre and CAR (n = 3). **(D).** Entropy of male short calls in Pre and CAR (n = 3). Signed-rank test.

**Supplementary Figure 3.**
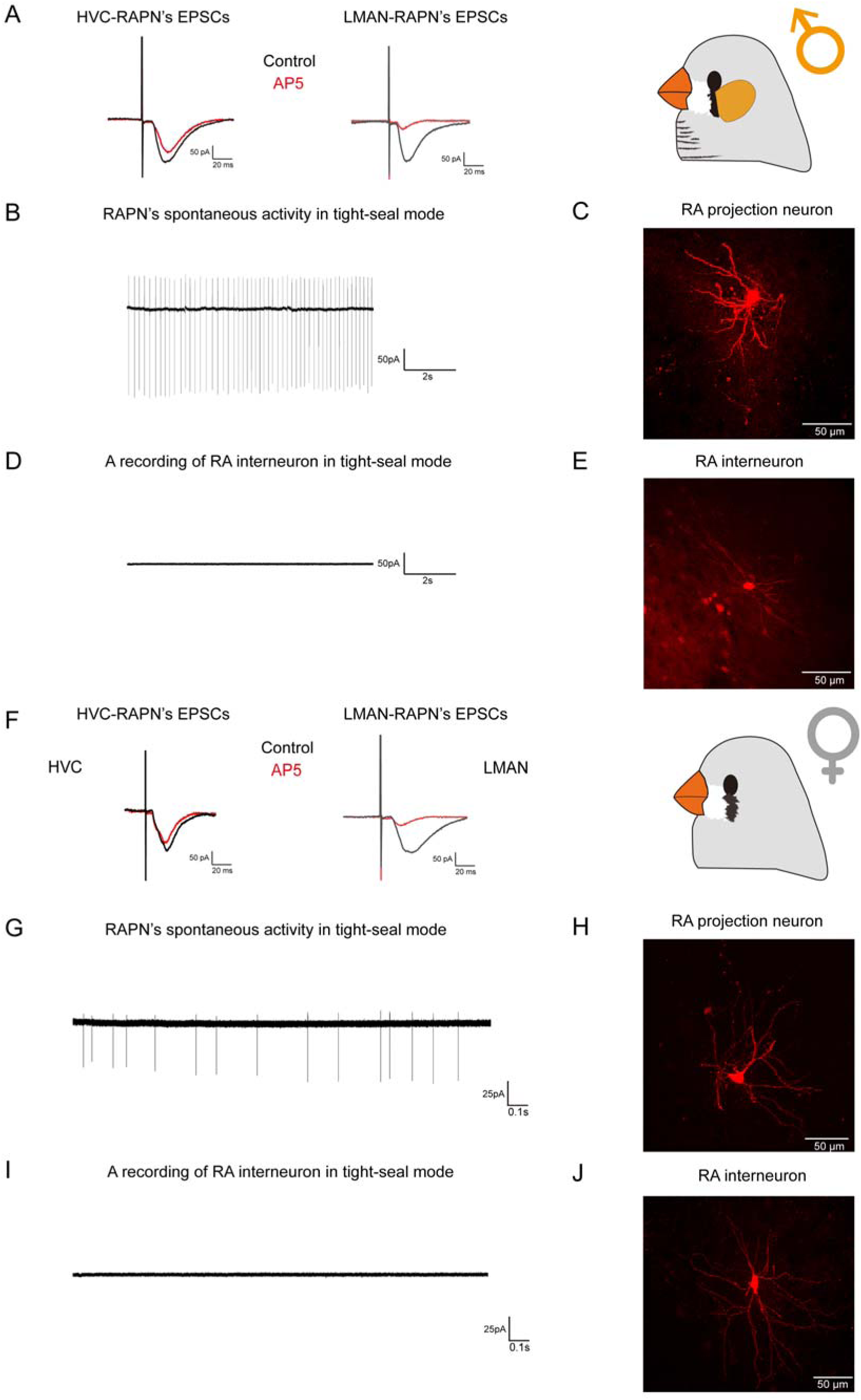
Pharmacological validation of HVC-RA and LMAN-RA pathways, as well as electrophysiological and morphological identification of RAPNs and RA interneurons. **(A).** In male adult zebra finches, HVC-RAPN’s EPSCs were weakly blocked by NMDA receptor antagonist AP5, while LMAN-RAPN’s EPSCs were mainly blocked by AP5. **(B).** A sample recording of male RAPNs’ spontaneous activities in tight-seal mode. **(C).** A fluorescence image of a male RAPN. **(D).** A sample recording of male RA interneurons in tight-seal mode. **(E).** A fluorescence image of a male RA interneuron. **(F).** Similar to (A), the glutamatergic pharmacology of EPSCs in female HVC-RAPNs and LMAN-RAPNs is the same as that in males. **(G).** A sample recording of female RAPNs’ spontaneous activities in tight-seal mode. **(H).** A fluorescence image of a female RAPN. **(I).** A sample recording of female RA interneurons in tight-seal mode. **(J).** A fluorescence image of a female RA interneuron.

**Supplementary Figure 4.**
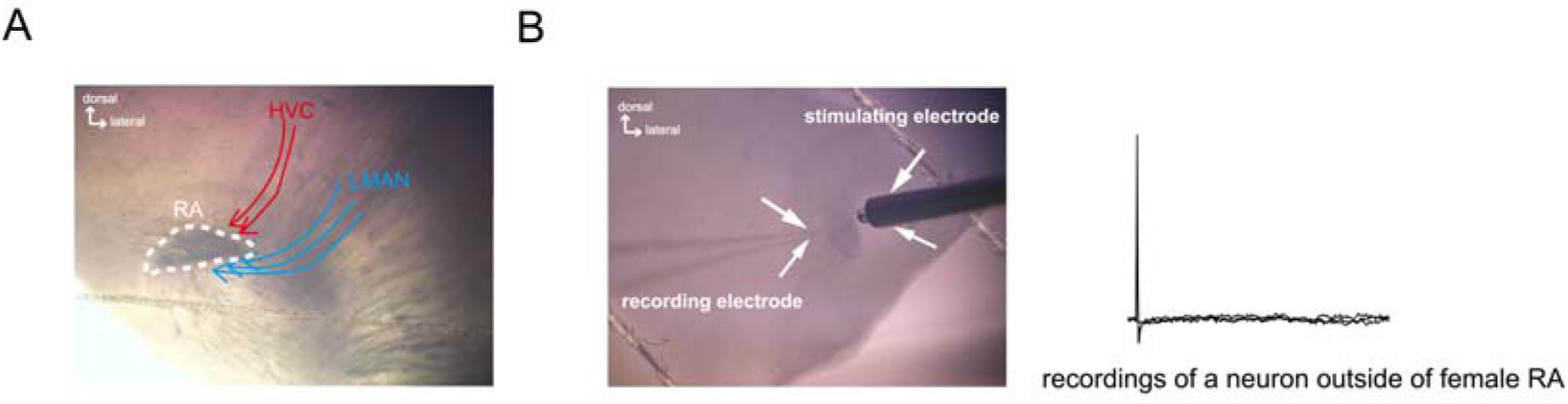
Female HVC-RA and LMAN-RA projections, and recordings of female RA peripheral neurons stimulated on LMAN-RA fibers. **(A).** An image of the coronal brain section of an adult female zebra finch displays female RA outline and the distribution of HVC-RA and LMAN-RA projections. **(B).** An example of recordings of a neuron outside of female RA showing no effect of stimulation on LMAN-RA fibers.

**Supplementary Figure 5.**
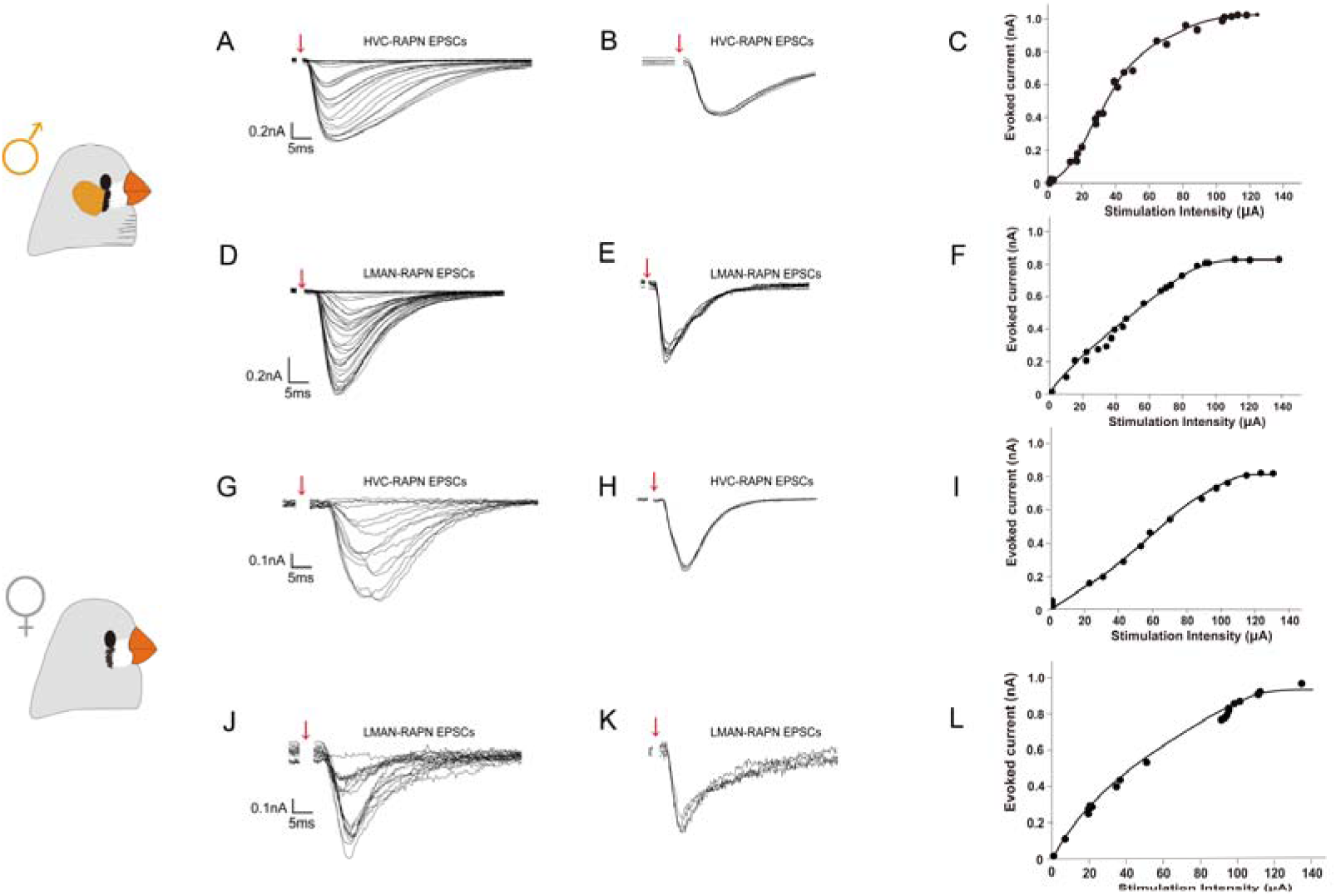
Examples of the stability test for EPSCs recordings show the reliable amplitudes of evoked EPSCs in both males and females. **(A).** An example of HVC-RAPN evoked EPSCs recordings for males shows that as the stimulation intensity gradually increased, the amplitude of EPSCs also increased until it reached its maximum. **(B).** An example of HVC-RAPN EPSCs recordings with stimulation every 1 min for males shows the reliable amplitudes of evoked EPSCs. **(C).** An example of the stimulation intensity/evoked current (I/O) curve of HVC-RAPN EPSCs recordings for males. **(D).** Similar to (A), an example of LMAN-RAPN evoked EPSCs recordings for males. **(E).** Similar to (B), an example of LMAN-RAPN EPSCs recordings with stimulation every 1 min for males. **(F).** An example of the I/O curve of LMAN-RAPN EPSCs recordings for males. **(G).** Similar to (A), an example of HVC-RAPN evoked EPSCs recordings for females. **(H).** Similar to (B), an example of HVC-RAPN EPSCs recordings with stimulation every 1 min for females. **(I).** An example of the I/O curve of HVC-RAPN EPSCs recordings for females. **(J).** Similar to (A), an example of LMAN-RAPN evoked EPSCs recordings for females. **(K).** Similar to (B), an example of LMAN-RAPN EPSCs recordings with stimulation every 1 min for females. **(L).** An example of the I/O curve of LMAN-RAPN EPSCs recordings for females.

**Supplementary Figure 6.**
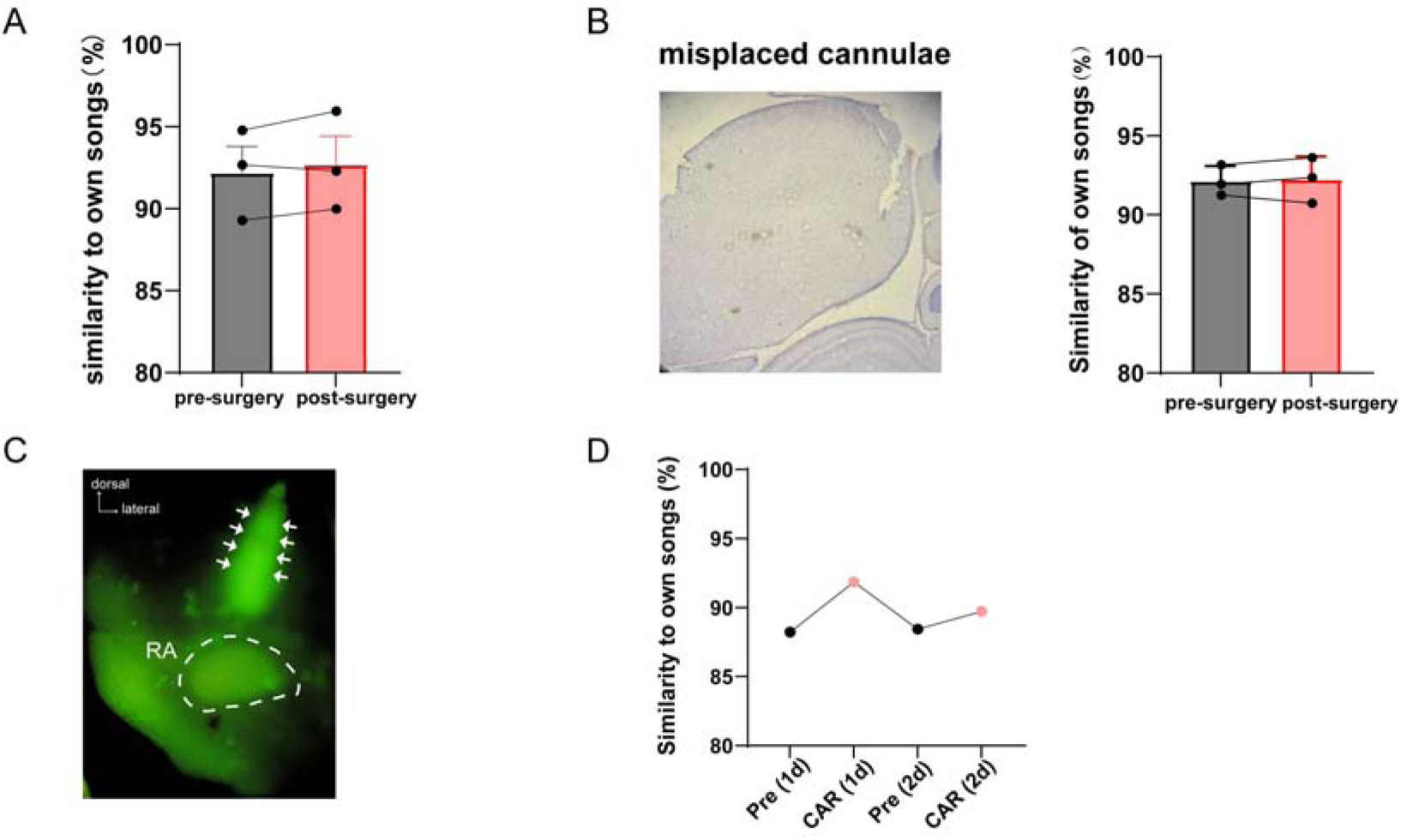
Validation of surgery impact, misplaced cannulas, drug diffusion and baseline before the 2nd administration. **(A).** The cannula implantation surgery did not have a significant impact on the song self-similarity (n = 3). Signed-rank test. **(B).** Left, a Nissl-stained single-side coronal section, which contains the cannula track but no RA, shows an example of the misplaced cannula. Right, the data of birds with misplaced cannulas show that misplaced cannulas did not have a significant impact on the song self-similarity, providing controls for the surgery and microinjection (n = 3). Signed-rank test. **(C).** Validation of the drug diffusion range using Alexa Fluor™ 488. The arrows denote the injection site. **(D).** Validation of song self-similarity baseline in a bird before the 2nd drug administration in the next day.

## References

1. Wood AN. New roles for dopamine in motor skill acquisition: lessons from primates, rodents, and songbirds. Journal of neurophysiology 125, 2361–2374 (2021).

2. Dayan E, Cohen LG. Neuroplasticity subserving motor skill learning. Neuron 72, 443–454 (2011).

3. Conner JM, Culberson A, Packowski C, Chiba AA, Tuszynski MH. Lesions of the Basal forebrain cholinergic system impair task acquisition and abolish cortical plasticity associated with motor skill learning. Neuron 38, 819–829 (2003).

4. Bariselli S, Mateo Y, Reuveni N, Lovinger DM. Gestational ethanol exposure impairs motor skills in female mice through dysregulated striatal dopamine and acetylcholine function. Neuropsychopharmacology: official publication of the American College of Neuropsychopharmacology 48, 1808–1820 (2023).

5. Zhang Y, Zhou L, Zuo J, Wang S, Meng W. Analogies of human speech and bird song: From vocal learning behavior to its neural basis. Frontiers in psychology 14, 1100969 (2023).

6. Yanagihara S, Yazaki-Sugiyama Y. Auditory experience-dependent cortical circuit shaping for memory formation in bird song learning. Nature communications 7, 11946 (2016).

7. Brainard MS, Doupe AJ. What songbirds teach us about learning. Nature 417, 351–358 (2002).

8. Friedrich SR, Nevue AA, Andrade ALP, Velho TAF, Mello CV. Emergence of sex-specific transcriptomes in a sexually dimorphic brain nucleus. Cell reports 40, 111152 (2022).

9. Nottebohm F, Stokes TM, Leonard CM. Central control of song in the canary, Serinus canarius. The Journal of comparative neurology 165, 457–486 (1976).

10. Vicario DS. Organization of the zebra finch song control system: II. Functional organization of outputs from nucleus Robustus archistriatalis. The Journal of comparative neurology 309, 486–494 (1991).

11. McDonald KS, Kirn JR. Anatomical plasticity in the adult zebra finch song system. The Journal of comparative neurology 520, 3673–3686 (2012).

12. Li R, Sakaguchi H. Cholinergic innervation of the song control nuclei by the ventral paleostriatum in the zebra finch: a double-labeling study with retrograde fluorescent tracers and choline acetyltransferase immunohistochemistry. Brain research 763, 239–246 (1997).

13. Reiner A, Perkel DJ, Bruce LL, Butler AB, Csillag A, Kuenzel W, Medina L, Paxinos G, Shimizu T, Striedter G, Wild M, Ball GF, Durand S, Güntürkün O, Lee DW, Mello CV, Powers A, White SA, Hough G, Kubikova L, Smulders TV, Wada K, Dugas-Ford J, Husband S, Yamamoto K, Yu J, Siang C, Jarvis ED; Avian Brain Nomenclature Forum. Revised nomenclature for avian telencephalon and some related brainstem nuclei. The Journal of comparative neurology 473, 377–414 (2004).

14. Sakaguchi H, Saito N. The acetylcholine and catecholamine contents in song control nuclei of zebra finch during song ontogeny. Brain research Developmental brain research 47, 313–317 (1989).

15. Asogwa NC, Mori C, Sánchez-Valpuesta M, Hayase S, Wada K. Inter– and intra-specific differences in muscarinic acetylcholine receptor expression in the neural pathways for vocal learning in songbirds. The Journal of comparative neurology 526, 2856–2869 (2018).

16. Asogwa NC, Toji N, He Z, Shao C, Shibata Y, Tatsumoto S, Ishikawa H, Go Y, Wada K. Nicotinic acetylcholine receptors in a songbird brain. The Journal of comparative neurology 530, 1966–1991 (2022).

17. Puzerey PA, Maher K, Prasad N, Goldberg JH. Vocal learning in songbirds requires cholinergic signaling in a motor cortex-like nucleus. Journal of neurophysiology 120, 1796–1806 (2018).

18. Yu AC, Margoliash D. Temporal hierarchical control of singing in birds. Science (New York, NY) 273, 1871–1875 (1996).

19. Leonardo A, Fee MS. Ensemble coding of vocal control in birdsong. The Journal of neuroscience: the official journal of the Society for Neuroscience 25, 652–661 (2005).

20. Hahnloser RH, Kozhevnikov AA, Fee MS. An ultra-sparse code underlies the generation of neural sequences in a songbird. Nature 419, 65–70 (2002).

21. Sober SJ, Wohlgemuth MJ, Brainard MS. Central contributions to acoustic variation in birdsong. The Journal of neuroscience: the official journal of the Society for Neuroscience 28, 10370–10379 (2008).

22. Meng W, Wang S, Yao L, Zhang N, Li D. Muscarinic Receptors Are Responsible for the Cholinergic Modulation of Projection Neurons in the Song Production Brain Nucleus RA of Zebra Finches. Frontiers in cellular neuroscience 11, 51 (2017).

23. Pérez SE, Yáñez J, Marín O, Anadón R, González A, Rodríguez-Moldes I. Distribution of choline acetyltransferase (ChAT) immunoreactivity in the brain of the adult trout and tract-tracing observations on the connections of the nuclei of the isthmus. The Journal of comparative neurology 428, 450–474 (2000).

24. Clemente D, Porteros A, Weruaga E, Alonso JR, Arenzana FJ, Aijón J, Arévalo R. Cholinergic elements in the zebrafish central nervous system: Histochemical and immunohistochemical analysis. The Journal of comparative neurology 474, 75–107 (2004).

25. López JM, Domínguez L, Morona R, Northcutt RG, González A. Organization of the cholinergic systems in the brain of two lungfishes, Protopterus dolloi and Neoceratodus forsteri. Brain structure & function 217, 549–576 (2012).

26. Hoogland PV, Vermeulen-VanderZee E. Distribution of choline acetyltransferase immunoreactivity in the telencephalon of the lizard Gekko gecko. Brain, behavior and evolution 36, 378–390 (1990).

27. Powers AS, Reiner A. The distribution of cholinergic neurons in the central nervous system of turtles. Brain, behavior and evolution 41, 326–345 (1993).

28. Pediconi MF, Roccamo de Fernández AM, Barrantes FJ. Asymmetric distribution and down-regulation of the muscarinic acetylcholine receptor in rat cerebral cortex. Neurochemical research 18, 565–572 (1993).

29. Tian MK, Bailey CD, Lambe EK. Cholinergic excitation in mouse primary vs. associative cortex: region-specific magnitude and receptor balance. The European journal of neuroscience 40, 2608–2618 (2014).

30. Ghimire M, Cai R, Ling L, Hackett TA, Caspary DM. Nicotinic Receptor Subunit Distribution in Auditory Cortex: Impact of Aging on Receptor Number and Function. The Journal of neuroscience: the official journal of the Society for Neuroscience 40, 5724–5739 (2020).

31. Benoy A, Bin Ibrahim MZ, Behnisch T, Sajikumar S. Metaplastic Reinforcement of Long-Term Potentiation in Hippocampal Area CA2 by Cholinergic Receptor Activation. The Journal of neuroscience: the official journal of the Society for Neuroscience 41, 9082–9098 (2021).

32. Medina L, Reiner A. Distribution of choline acetyltransferase immunoreactivity in the pigeon brain. The Journal of comparative neurology 342, 497–537 (1994).

33. Dietl MM, Cortés R, Palacios JM. Neurotransmitter receptors in the avian brain. II. Muscarinic cholinergic receptors. Brain research 439, 360–365 (1988).

34. Lohmann TH, Torrão AS, Britto LR, Lindstrom J, Hamassaki-Britto DE. A comparative non-radioactive in situ hybridization and immunohistochemical study of the distribution of alpha7 and alpha8 subunits of the nicotinic acetylcholine receptors in visual areas of the chick brain. Brain research 852, 463–469 (2000).

35. Semba K. Phylogenetic and ontogenetic aspects of the basal forebrain cholinergic neurons and their innervation of the cerebral cortex. Progress in brain research 145, 3–43 (2004).

36. Ztaou S, Maurice N, Camon J, Guiraudie-Capraz G, Kerkerian-Le Goff L, Beurrier C, Liberge M, Amalric M. Involvement of Striatal Cholinergic Interneurons and M1 and M4 Muscarinic Receptors in Motor Symptoms of Parkinson’s Disease. The Journal of neuroscience: the official journal of the Society for Neuroscience 36, 9161–9172 (2016).

37. Power AE, Vazdarjanova A, McGaugh JL. Muscarinic cholinergic influences in memory consolidation. Neurobiology of learning and memory 80, 178–193 (2003).

38. Hasselmo ME. The role of acetylcholine in learning and memory. Current opinion in neurobiology 16, 710–715 (2006).

39. Greco MA, McCarley RW, Shiromani PJ. Choline acetyltransferase expression during periods of behavioral activity and across natural sleep-wake states in the basal forebrain. Neuroscience 93, 1369–1374 (1999).

40. Nikonova EV, Gilliland JD, Tanis KQ, Podtelezhnikov AA, Rigby AM, Galante RJ, Finney EM, Stone DJ, Renger JJ, Pack AI, Winrow CJ. Transcriptional Profiling of Cholinergic Neurons From Basal Forebrain Identifies Changes in Expression of Genes Between Sleep and Wake. Sleep 40, (2017).

41. Sarter M, Lustig C, Blakely RD, Koshy Cherian A. Cholinergic genetics of visual attention: Human and mouse choline transporter capacity variants influence distractibility. Journal of physiology, Paris 110, 10–18 (2016).

42. Orciani C, Hall H, Pentz R, Foret MK, Do Carmo S, Cuello AC. Long-term nucleus basalis cholinergic depletion induces attentional deficits and impacts cortical neurons and BDNF levels without affecting the NGF synthesis. Journal of neurochemistry 163, 149–167 (2022).

43. Williams VM, Bhagwandin A, Swiegers J, Bertelsen MF, Hård T, Sherwood CC, Manger PR. Distribution of cholinergic neurons in the brains of a lar gibbon and a chimpanzee. Anatomical record (Hoboken, NJ: 2007) 305, 1516–1535 (2022).

44. Yang D, Günter R, Qi G, Radnikow G, Feldmeyer D. Muscarinic and Nicotinic Modulation of Neocortical Layer 6A Synaptic Microcircuits Is Cooperative and Cell-Specific. Cerebral cortex (New York, NY: 1991) 30, 3528–3542 (2020).

45. Roberts TF, Hall WS, Brauth SE. Organization of the avian basal forebrain: chemical anatomy in the parrot (Melopsittacus undulatus). The Journal of comparative neurology 454, 383–408 (2002).

46. Ryan SM, Arnold AP. Evidence for cholinergic participation in the control of bird song; acetylcholinesterase distribution and muscarinic receptor autoradiography in the zebra finch brain. The Journal of comparative neurology 202, 211–219 (1981).

47. Sadananda M. Acetylcholinesterase in central vocal control nuclei of the zebra finch (Taeniopygia guttata). Journal of biosciences 29, 189–200 (2004).

48. Zuschratter W, Scheich H. Distribution of choline acetyltransferase and acetylcholinesterase in the vocal motor system of zebra finches. Brain research 513, 193–201 (1990).

49. Lovell PV, Huizinga NA, Friedrich SR, Wirthlin M, Mello CV. The constitutive differential transcriptome of a brain circuit for vocal learning. BMC genomics 19, 231 (2018).

50. Lovell PV, Clayton DF, Replogle KL, Mello CV. Birdsong “transcriptomics”: neurochemical specializations of the oscine song system. PloS one 3, e3440 (2008).

51. Watson JT, Adkins-Regan E, Whiting P, Lindstrom JM, Podleski TR. Autoradiographic localization of nicotinic acetylcholine receptors in the brain of the zebra finch (Poephila guttata). The Journal of comparative neurology 274, 255–264 (1988).

52. Conner JM, Kulczycki M, Tuszynski MH. Unique contributions of distinct cholinergic projections to motor cortical plasticity and learning. Cerebral cortex (New York, NY: 1991) 20, 2739–2748 (2010).

53. Li Y, Hollis E. Basal Forebrain Cholinergic Neurons Selectively Drive Coordinated Motor Learning in Mice. The Journal of neuroscience: the official journal of the Society for Neuroscience 41, 10148–10160 (2021).

54. Voegtle A, Mohrbutter C, Hils J, Schulz S, Weuthen A, Brämer U, Ullsperger M, Sweeney-Reed CM. Cholinergic modulation of motor sequence learning. The European journal of neuroscience, (2024).

55. Cappendijk SL, Pirvan DF, Miller GL, Rodriguez MI, Chalise P, Halquist MS, James JR. In vivo nicotine exposure in the zebra finch: a promising innovative animal model to use in neurodegenerative disorders related research. Pharmacology, biochemistry, and behavior 96, 152–159 (2010).

56. Perry WM, Cappendijk SL. Effects of nicotine administration on spectral and temporal features of crystallized song in the adult male zebra finch. Nicotine & tobacco research: official journal of the Society for Research on Nicotine and Tobacco 16, 1409–1416 (2014).

57. Chi Z, Margoliash D. Temporal precision and temporal drift in brain and behavior of zebra finch song. Neuron 32, 899–910 (2001).

58. Tian LY, Warren TL, Mehaffey WH, Brainard MS. Dynamic top-down biasing implements rapid adaptive changes to individual movements. eLife 12, (2023).

59. Shea SD, Margoliash D. Basal forebrain cholinergic modulation of auditory activity in the zebra finch song system. Neuron 40, 1213–1226 (2003).

60. Shea SD, Koch H, Baleckaitis D, Ramirez JM, Margoliash D. Neuron-specific cholinergic modulation of a forebrain song control nucleus. Journal of neurophysiology 103, 733–745 (2010).

61. Jaffe PI, Brainard MS. Acetylcholine acts on songbird premotor circuitry to invigorate vocal output. eLife 9, (2020).

62. Sakaguchi H, Saito N. Developmental change of cholinergic activity in the forebrain of the zebra finch during song learning. Brain research Developmental brain research 62, 223–228 (1991).

63. Sakaguchi H. Developmental changes in carbachol-stimulated phosphoinositide turnover in synaptoneurosomes of the robust nucleus of the archistriatum in the zebra finch. Neuroreport 6, 1901–1904 (1995).

64. Salgado-Commissariat D, Rosenfield DB, Helekar SA. Nicotine-mediated plasticity in robust nucleus of the archistriatum of the adult zebra finch. Brain research 1018, 97–105 (2004).

65. Pfenning AR, Hara E, Whitney O, Rivas MV, Wang R, Roulhac PL, Howard JT, Wirthlin M, Lovell PV, Ganapathy G, Mouncastle J, Moseley MA, Thompson JW, Soderblom EJ, Iriki A, Kato M, Gilbert MT, Zhang G, Bakken T, Bongaarts A, Bernard A, Lein E, Mello CV, Hartemink AJ, Jarvis ED. Convergent transcriptional specializations in the brains of humans and song-learning birds. Science (New York, NY) 346, 1256846 (2014).

66. Jarvis ED. Evolution of vocal learning and spoken language. Science (New York, NY) 366, 50–54 (2019).

67. Meng W, Wang SH, Li DF. Carbachol-Induced Reduction in the Activity of Adult Male Zebra Finch RA Projection Neurons. Neural plasticity 2016, 7246827 (2016).

68. Levy RB, Reyes AD, Aoki C. Nicotinic and muscarinic reduction of unitary excitatory postsynaptic potentials in sensory cortex; dual intracellular recording in vitro. Journal of neurophysiology 95, 2155–2166 (2006).

69. McCasland JS. Neuronal control of bird song production. The Journal of neuroscience: the official journal of the Society for Neuroscience 7, 23–39 (1987).

70. Bottjer SW, Brady JD, Cribbs B. Connections of a motor cortical region in zebra finches: relation to pathways for vocal learning. The Journal of comparative neurology 420, 244–260 (2000).

71. Glasgow SD, McPhedrain R, Madranges JF, Kennedy TE, Ruthazer ES. Approaches and Limitations in the Investigation of Synaptic Transmission and Plasticity. Frontiers in synaptic neuroscience 11, 20 (2019).

72. Tryon SC, Bratsch-Prince JX, Warren JW, Jones GC, McDonald AJ, Mott DD. Differential Regulation of Prelimbic and Thalamic Transmission to the Basolateral Amygdala by Acetylcholine Receptors. The Journal of neuroscience: the official journal of the Society for Neuroscience 43, 722–735 (2023).

73. Spiro JE, Dalva MB, Mooney R. Long-range inhibition within the zebra finch song nucleus RA can coordinate the firing of multiple projection neurons. Journal of neurophysiology 81, 3007–3020 (1999).

74. Zemel BM, Nevue AA, Tavares LES, Dagostin A, Lovell PV, Jin DZ, Mello CV, von Gersdorff H. Motor cortex analogue neurons in songbirds utilize Kv3 channels to generate ultranarrow spikes. eLife 12, (2023).

75. Wood WE, Lovell PV, Mello CV, Perkel DJ. Serotonin, via HTR2 receptors, excites neurons in a cortical-like premotor nucleus necessary for song learning and production. The Journal of neuroscience: the official journal of the Society for Neuroscience 31, 13808–13815 (2011).

76. Wood WE, Roseberry TK, Perkel DJ. HTR2 receptors in a songbird premotor cortical-like area modulate spectral characteristics of zebra finch song. The Journal of neuroscience: the official journal of the Society for Neuroscience 33, 2908–2915 (2013).

77. Wang S, Liu S, Wang Q, Sun Y, Yao L, Li D, Meng W. Dopamine Modulates Excitatory Synaptic Transmission by Activating Presynaptic D1-like Dopamine Receptors in the RA Projection Neurons of Zebra Finches. Frontiers in cellular neuroscience 14, 126 (2020).

78. Solis MM, Perkel DJ. Noradrenergic modulation of activity in a vocal control nucleus in vitro. Journal of neurophysiology 95, 2265–2276 (2006).

79. Sizemore M, Perkel DJ. Noradrenergic and GABA B receptor activation differentially modulate inputs to the premotor nucleus RA in zebra finches. Journal of neurophysiology 100, 8–18 (2008).

80. Wang S, Sun Y, Wang Q, Qiu Y, Yao L, Gong Y, Meng W, Li D. Sexual dimorphism of inhibitory synaptic transmission in RA projection neurons of songbirds. Neuroscience letters 709, 134377 (2019).

81. Scharff C, Nottebohm FA comparative study of the behavioral deficits following lesions of various parts of the zebra finch song system: implications for vocal learning. The Journal of neuroscience: the official journal of the Society for Neuroscience 11, 2896–2913 (1991).

82. Warren TL, Tumer EC, Charlesworth JD, Brainard MS. Mechanisms and time course of vocal learning and consolidation in the adult songbird. Journal of neurophysiology 106, 1806–1821 (2011).

83. Kao MH, Doupe AJ, Brainard MS. Contributions of an avian basal ganglia-forebrain circuit to real-time modulation of song. Nature 433, 638–643 (2005).

84. Andalman AS, Fee MS. A basal ganglia-forebrain circuit in the songbird biases motor output to avoid vocal errors. Proceedings of the National Academy of Sciences of the United States of America 106, 12518–12523 (2009).

85. Aronov D, Andalman AS, Fee MS. A specialized forebrain circuit for vocal babbling in the juvenile songbird. *Science (New York*, NY) 320, 630–634 (2008).

86. Bottjer SW, Miesner EA, Arnold AP. Forebrain Lesions Disrupt Development But Not Maintenance of Song in Passerine Birds. *Science (New York*, NY) 224, 901–903 (1984).

87. Herrmann K, Arnold AP. The development of afferent projections to the robust archistriatal nucleus in male zebra finches: a quantitative electron microscopic study. The Journal of neuroscience: the official journal of the Society for Neuroscience 11, 2063–2074 (1991).

88. Brainard MS. Contributions of the anterior forebrain pathway to vocal plasticity. Annals of the New York Academy of Sciences 1016, 377–394 (2004).

89. Nordeen KW, Nordeen EJ. Deafening-induced vocal deterioration in adult songbirds is reversed by disrupting a basal ganglia-forebrain circuit. The Journal of neuroscience: the official journal of the Society for Neuroscience 30, 7392–7400 (2010).

90. Chen Z, Ye R, Goldman SA. Testosterone modulation of angiogenesis and neurogenesis in the adult songbird brain. Neuroscience 239, 139–148 (2013).

91. Nottebohm F, Arnold AP. Sexual dimorphism in vocal control areas of the songbird brain. *Science (New York*, NY) 194, 211–213 (1976).

92. Brenowitz EA. Testosterone and brain-derived neurotrophic factor interactions in the avian song control system. Neuroscience 239, 115–123 (2013).

93. Wade J, Arnold AP. Sexual differentiation of the zebra finch song system. Annals of the New York Academy of Sciences 1016, 540–559 (2004).

94. Liu XL, Hou GQ, Liao SQ, Li DF. Sexual dimorphism of the electrophysiological properties of the projection neurons in the robust nucleus of the arcopallium in adult zebra finches. Neuroscience bulletin 26, 147–152 (2010).

95. Zemel BM, Nevue AA, Dagostin A, Lovell PV, Mello CV, von Gersdorff H. Resurgent Na(+) currents promote ultrafast spiking in projection neurons that drive fine motor control. Nature communications 12, 6762 (2021).

96. Wang S, Meng W, Liu S, Liao C, Huang Q, Li D. Sex differences of excitatory synaptic transmission in RA projection neurons of adult zebra finches. Neuroscience letters 582, 75–80 (2014).

97. Bottjer SW, Roselinsky H, Tran NB. Sex differences in neuropeptide staining of song-control nuclei in zebra finch brains. Brain, behavior and evolution 50, 284–303 (1997).

98. Sakaguchi H, Li R, Taniguchi I. Sex differences in the ventral paleostriatum of the zebra finch: origin of the cholinergic innervation of the song control nuclei. Neuroreport 11, 2727–2731 (2000).

99. Konishi M, Akutagawa E. Neuronal growth, atrophy and death in a sexually dimorphic song nucleus in the zebra finch brain. Nature 315, 145–147 (1985).

100. Mooney R, Rao M. Waiting periods versus early innervation: the development of axonal connections in the zebra finch song system. The Journal of neuroscience: the official journal of the Society for Neuroscience 14, 6532–6543 (1994).

101. Holloway CC, Clayton DF. Estrogen synthesis in the male brain triggers development of the avian song control pathway in vitro. Nature neuroscience 4, 170–175 (2001).

102. Diez A, Wang S, Carfagnini N, MacDougall-Shackleton SA. Sex differences in myelination of the zebra finch vocal control system emerge relatively late in development. Developmental neurobiology 82, 581–595 (2022).

103. Johnson F, Sellix M. Reorganization of a telencephalic motor region during sexual differentiation and vocal learning in zebra finches. Brain research Developmental brain research 121, 253–263 (2000).

104. Shaughnessy DW, Hyson RL, Bertram R, Wu W, Johnson F. Female zebra finches do not sing yet share neural pathways necessary for singing in males. The Journal of comparative neurology 527, 843–855 (2019).

105. Wang J, Sakaguchi H, Sokabe M. Sex differences in the vocal motor pathway of the zebra finch revealed by real-time optical imaging technique. Neuroreport 10, 2487–2491 (1999).

106. Nixdorf-Bergweiler BE. Divergent and parallel development in volume sizes of telencephalic song nuclei in male and female zebra finches. The Journal of comparative neurology 375, 445–456 (1996).

107. Nixdorf-Bergweiler BE. Lateral magnocellular nucleus of the anterior neostriatum (LMAN) in the zebra finch: neuronal connectivity and the emergence of sex differences in cell morphology. Microscopy research and technique 54, 335–353 (2001).

108. Nordeen EJ, Grace A, Burek MJ, Nordeen KW. Sex-dependent loss of projection neurons involved in avian song learning. Journal of neurobiology 23, 671–679 (1992).

109. Hamilton KS, King AP, Sengelaub DR, West MJ. A brain of her own: a neural correlate of song assessment in a female songbird. Neurobiology of learning and memory 68, 325–332 (1997).

110. Riebel K, Smallegange IM, Terpstra NJ, Bolhuis JJ. Sexual equality in zebra finch song preference: evidence for a dissociation between song recognition and production learning. Proceedings Biological sciences 269, 729–733 (2002).

111. Terpstra NJ, Bolhuis JJ, Riebel K, van der Burg JM, den Boer-Visser AM. Localized brain activation specific to auditory memory in a female songbird. The Journal of comparative neurology 494, 784–791 (2006).

112. Zhang Y, Wang Q, Zheng Z, Sun Y, Niu Y, Li D, Wang S, Meng W. BDNF enhances electrophysiological activity and excitatory synaptic transmission of RA projection neurons in adult male zebra finches. Brain research 1801, 148208 (2023).

113. Mehaffey WH, Doupe AJ. Naturalistic stimulation drives opposing heterosynaptic plasticity at two inputs to songbird cortex. Nature neuroscience 18, 1272–1280 (2015).

114. Chung JH, Bottjer SW. Developmentally regulated pathways for motor skill learning in songbirds. The Journal of comparative neurology 530, 1288–1301 (2022).

115. Garst-Orozco J, Babadi B, Ölveczky BP. A neural circuit mechanism for regulating vocal variability during song learning in zebra finches. eLife 3, e03697 (2014).

116. Simpson HB, Vicario DS. Brain pathways for learned and unlearned vocalizations differ in zebra finches. The Journal of neuroscience: the official journal of the Society for Neuroscience 10, 1541–1556 (1990).

117. Tchernichovski O, Nottebohm F, Ho CE, Pesaran B, Mitra PP. A procedure for an automated measurement of song similarity. Animal behaviour 59, 1167–1176 (2000).

